# Prediction of inhibitory peptides against *E. coli* with desired MIC value

**DOI:** 10.1101/2024.07.18.604028

**Authors:** Nisha Bajiya, Nishant Kumar, Gajendra P. S. Raghava

**Affiliations:** Department of Computational Biology, Indraprastha Institute of Information Technology, Okhla Phase 3, New Delhi-110020, India

**Keywords:** Inhibitory peptides, Minimum Inhibitory Concentration, Machine Learning, *Escherichia coli*, Peptide Design, Regression models

## Abstract

In the past, several methods have been developed for predicting antibacterial and antimicrobial peptides, but only limited attempts have been made to predict their minimum inhibitory concentration (MIC) values. In this study, we trained our models on 3,143 peptides and validated them on 786 peptides whose MIC values have been determined experimentally against *Escherichia coli* (*E. coli*). The correlational analysis reveals that the Composition Enhanced Transition and Distribution (CeTD) attributes strongly correlate with MIC values. We initially employed the similarity search strategy utilizing BLAST to estimate MIC values of peptides but found it inadequate for prediction. Next, we developed machine learning techniques-based regression models using a wide range of features, including peptide composition, binary profile, and embeddings of large language models. We implemented feature selection techniques like minimum Redundancy Maximum Relevance (mRMR) to select the best relevant features for developing prediction models. Our Random forest-based regressor, based on selected features, achieved a correlation coefficient (R) of 0.78, R-squared (R²) of 0.59, and a root mean squared error (RMSE) of 0.53 on the validation dataset. Our best model outperforms the existing methods when benchmarked on an independent dataset of 498 inhibitory peptides of *E. coli*. One of the major features of the web-based platform EIPpred developed in this study is that it allows users to identify or design peptides that can inhibit *E. coli* with the desired MIC value (https://webs.iiitd.edu.in/raghava/eippred).

**Highlights:**

- Prediction of MIC value of peptides against *E.coli*.
- An independent dataset was generated for comparison.
- Feature selection using the mRMR method.
- A regressor method for designing novel inhibitory peptides.
- A web server and standalone package for predicting the inhibitory activity of peptides.

## 1. Introduction

The introduction of antibiotics in the 1940s revolutionized medicine by effectively treating bacterial infections and reducing mortality rates. However, issues such as antibiotic resistance and allergic reactions quickly surfaced[1]. By the early 21st century, resistance had rendered many antibiotics ineffective, causing an estimated 5 million deaths annually, with projections rising to 10 million by 2050 [2]. The escalating resistance of pathogens to conventional antibiotics has become a global crisis, demanding new treatment approaches. In response, alternative therapies like biologics-based treatments have garnered interest for their specificity and reduced potential for resistance development [3]. These innovative strategies offer a promising solution to the limitations of traditional antibiotics in the current era of resistance. Among the leading alternatives are peptide/protein-based therapies, including antimicrobial peptides (AMPs) and monoclonal antibodies (mAbs). AMPs are highly effective against a range of pathogens, including multidrug-resistant ones, due to their rapid action and unique mechanisms that limit resistance development. Part of the innate immune system, AMPs are generally amphipathic and contain cationic amino acids, which enable them to disrupt the negatively charged bacterial membrane [4]. They work through mechanisms such as membrane disruption (carpet formation, barrel stave formation, toroidal pore formation) or non-membrane pathways, including the inhibition of protein synthesis, DNA/RNA synthesis, metabolic processes, or cell wall formation [Figure 1] [5].

**Figure 1:**
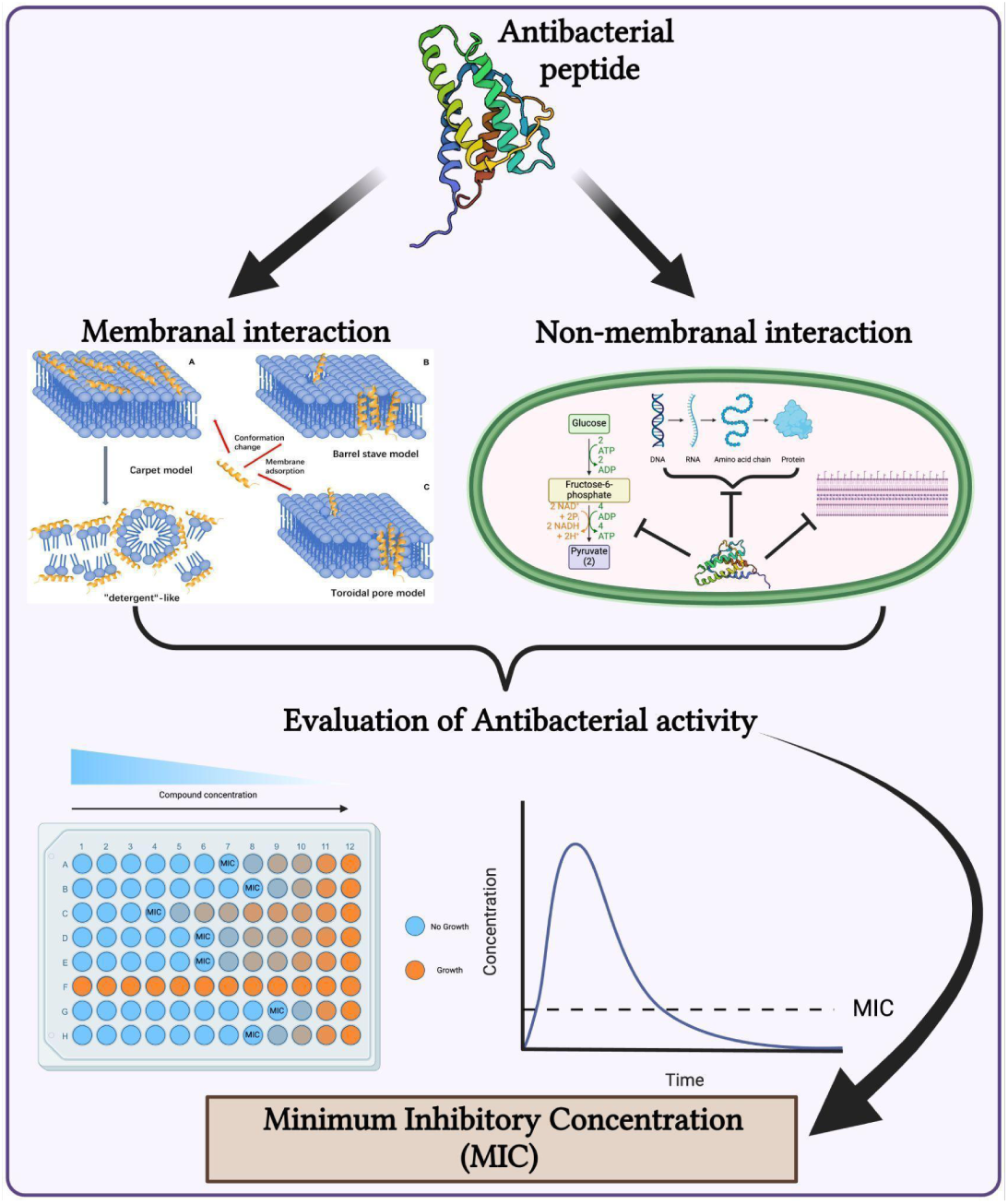
Graphical representation of the working of antibacterial peptides and their activity prediction.

In the field of peptides, extensive research has led to the creation of comprehensive databases accessible to the scientific community. The first such database, the Antimicrobial Peptide Database (APD), was established in 2004 and contains detailed information on peptides with various properties, including antitumor, antiviral, antifungal, and antibacterial activities [6]. Since its inception, numerous public repositories have emerged, providing data on natural, synthetic, and predicted antimicrobial/antibacterial peptides. Notable examples include CAMPR4, DBAASP v3, DRAMP 3.0 and AntiTbPdb, which offer comprehensive information on the peptide structure, functional activities, target species, and sources[7][8][9][10]. These databases, populated with data from both experimental methods and computational resources, are invaluable for designing new drugs and advancing peptide research.

With the growth of publicly available data on AMPs, there has been a significant increase in the development of accurate prediction tools for identifying and designing AMP sequences. Examples of these methods include AI4AMP, AMPDiscover, AMPScanner vr.2, AntiBP3, AntiFP and AntiMPmod [11][12][13][14][15][16]. Similarly, species-specific predictive methods like AntiTbPred are designed to distinguish antituberculosis peptides[17]. These methods aim to predict peptides that can inhibit bacterial growth but do not assess the inhibitory potential or the amount of peptide required to inhibit bacterial growth [18]. Recent studies have attempted to predict the minimum inhibitory concentration (MIC) value of peptides against *Escherichia coli* (*E. coli*) or other bacteria [5][19][20][21][22]. To complement these existing methods, we have systematically developed an improved method for predicting the MIC value of peptides against *E. coli*. One of the major objectives of our study is to facilitate the scientific community for designing peptides with desired MIC values against *E. coli*. Therefore, we developed standalone software called EIPpred and a web-based platform to support this goal. Our standalone server can be used to scan antibacterial peptides in proteins against *E. coli* at a proteome scale. In addition to prediction, our server maps residues in peptides responsible for increasing or decreasing MIC values.

## 2. Materials and Methods

### 2.1. Collection of Dataset

#### Main dataset

The pre-processed dataset of 3929 peptides with MIC value against *E. coli* was obtained from paper MBC-CNN attention, which was originally curated from the DBAASP v3 database [23][8]. This dataset was divided into training and validation sets, with the training set containing 3,143 (80%) sequences and the validation set containing 786 (20%) sequences. To avoid biases in peptide lengths, we ensured that the average length was the same in both the training and validation datasets.

#### Independent dataset

We have already used 20% of the data to validate our models using external validation. In addition, we have created an independent dataset of peptides with MIC values against *E. coli* to evaluate the performance of our final model and the existing methods. We extracted 498 unique experimentally validated peptides, curated from DBAASP [8], DRAMP [24] and APD3 [6] databases with a count of 127, 342 and 29 peptides, respectively. We ensured none of these sequences were identical to the training and validation dataset. We removed all those 8-60 length peptides that contain unnatural amino acids, ‘BJOUZX’.

#### Target variable

In this study, the target MIC value is represented as negative logarithmic MIC (in microMolar) against the bacteria, represented in equation 1. However, a single peptide can have multiple MIC values; in that scenario, the mean of all MIC values is considered the target prediction value for such multiple entries [21].

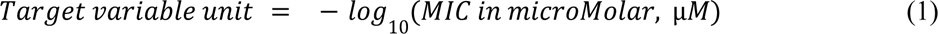

### 2.2. Workflow of the study

The detailed workflow of the study is represented in Figure 2, which depicts the progression of this study from the data curation to the final tool development and web server development.

**Figure 2:**
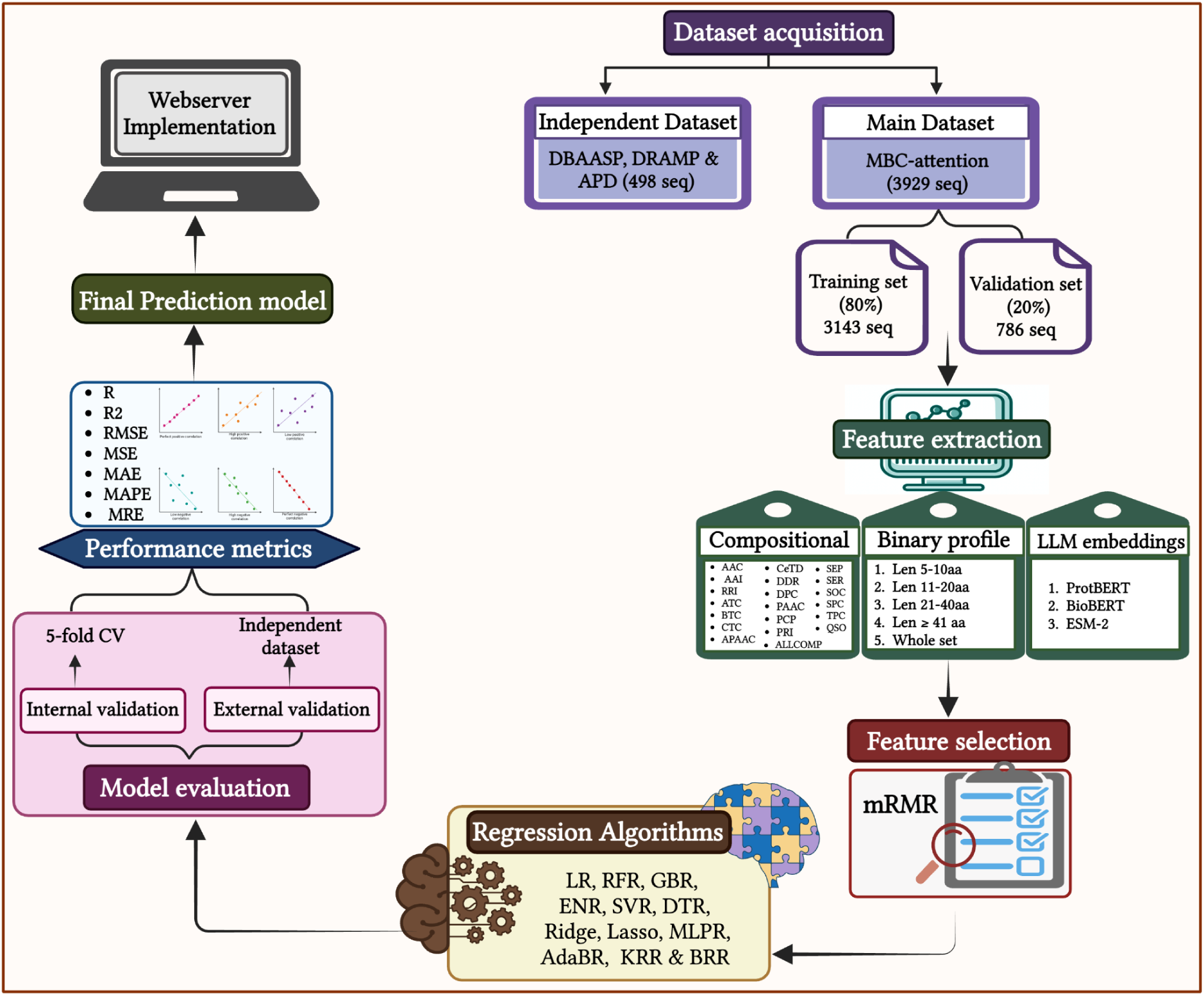
The detailed workflow of the study.

### 2.3. Cross-Validation

To ensure accurate error estimation, we employed a 5-fold cross-validation (CV) approach, which prevents overfitting, reduces bias, and allows us to evaluate the predictive ability of our models. We applied the 5-fold CV technique on the training dataset while keeping the validation dataset untouched. This approach divides the entire training dataset into five parts: four parts for training the model and the fifth part for testing it. This process is repeated five times, with each part serving as a test set once. The overall performance is calculated as the average of these five iterations. The untouched validation dataset is then used to externally validate our models.

### 2.4. BLAST for similarity search

BLAST is heavily used in literature to annotate protein sequences [25][26][27]. We have used it to identify peptides with antibacterial activity based on the similarity of peptides with known antibacterial activity, believing that similar sequences might have the same functional properties. The blastp (protein-protein BLAST) suite of BLAST+ version 2.7.1 was used to develop the similarity-based search module [28], where the query sequences were hit against the custom database built using the training set consisting of known activity of antibacterial peptides (ABPs). Then, a peptide is searched against a custom database using BLAST for different E-value cutoffs. For the hits obtained after running the BLAST, the validation peptide was assigned to the particular MIC values from the database, and the performance was evaluated using various e-values.

### 2.5. Peptide features

#### 2.5.1. Composition-based features

Residue information of the protein was used in the form of various compositional features for developing ML regression models. With the intent to calculate a diverse range of features from the sequences of protein or peptide, we have implemented the Pfeature package[29]. We deployed the composition-based module of Pfeature to generate more than 9000 descriptors of peptide sequences in the datasets. We have calculated nineteen types of features (AAC, DPC, RRI, DDR, SEP, SER, SPC, PRI, AAI, CTC, CeTD, PAAC, APAAC, QSO, TPC, ATC, PCP, BTC and SOC) as shown in Table 1. The input vector of 9189 descriptors was used further for feature selection and ML purposes[30].

**Table 1:** List of compositional features used in this study.

| S.No. | Acronym | Descriptors/Features | Number |
| --- | --- | --- | --- |
| 1 | AAC | Amino acid Composition | 20 |
| 2 | AAI | Amino Acid Index | 553 |
| 3 | APAAC | Amphiphilic Pseudo Amino Acid Composition | 23 |
| 4 | ATC | Atomic Composition | 5 |
| 5 | BTC | Bond Composition | 4 |
| 6 | CTC | Conjoint Triad Descriptors | 343 |
| 7 | CeTD | Composition-enhanced Transition Distribution | 189 |
| 8 | DDR | Distance Distribution of Residues | 20 |
| 9 | DPC | Dipeptide Composition | 400 |
| 10 | PAAC | Pseudo Amino Acid Composition | 21 |
| 11 | PCP | Physico-chemical properties | 19 |
| 12 | PRI | Property Repeats Index | 24 |
| 13 | QSO | Quasi-Sequence Order | 42 |
| 14 | RRI | Repetitive Residue Information | 20 |
| 15 | SEP | Shannon Entropy at Protein Level | 1 |
| 16 | SER | Shannon Entropy at Residue Level | 20 |
| 17 | SOC | Sequence Order Coupling number | 2 |
| 18 | SPC | Shannon Entropy at Property Level | 25 |
| 19 | TPC | Tripeptide Composition | 8000 |
| 20 | ALLCOMP | All Compositions | 9189 |

#### 2.5.2. Binary Profiles

We generated a binary profile or one hot encoding of patterns by assigning binary values to the amino acids in fixed-length patterns. First, we have divided the whole data set into four different groups according to the length of the peptides: 1st group with a length of 5-10aa, 2nd group with a length of 11-20aa, 3rd group with a length of 21-40aa and 4th group with a length of 41aa or more. For each group, we have further created fixed length vectors such as for group 5-10aa, a vector of 10 residues is formed by taking five residues from the N-terminal and five residues from the C-terminal, then generating a pattern with a vector size of 10×20 (here, 20 corresponds to each amino acid present in the pattern and 10 is the length of the pattern) [31]. Similarly, patterns of 20×20, 40×20 and 80×20 are generated for the second, third and fourth groups, respectively, as depicted in Supplementary Table S1.

#### 2.5.3. Large Language Model (LLM) embeddings

Transformer-based language models have recently been prominent in natural language processing (NLP) and have been applied to protein sequences. These protein language models (PLMs) seek to enhance protein representation by learning contextualized representation (also called embeddings) from an abundance of protein sequence data, improving tasks involving protein functions[32].

We evaluated three pre-trained protein models (protBERT, BioBERT and ESM-2) to extract features from the peptide sequences for regressive analysis. ProtBert [33] was trained on the Big Fantastic Database (BFD), which contains over 2.3 million protein sequences and is based on the BERT algorithm [34] [35]. The last attention layer produces an output of a 1024-dimensional embedding for each residue [36]. BioBERT (Bidirectional Encoder Representations from Transformers for Biomedical Text Mining) is a model pre-trained on large-scale biomedical corpora (PubMed abstracts and PubMed Central full-text articles) [37] that generates a total of 768 embeddings and ESM-2 (Evolutionary Scale Modeling) [38] is a general-purpose PLM based on the BERT transformer architecture and trained on UniRef50 to predict masked amino acids using all the preceding and following amino acids in the sequence. In this study, we used a model called esm2_t33_650M_UR50D (termed ESM-2), which has approximately 3 billion learnable parameters. The model’s output was an embedding of a feature dimension of 1280 for each amino acid [32].

### 2.6. Feature selection

Pfeature has generated a large number of peptide features, i.e., 9189, and all might not be relevant; some may be highly correlated with each other and can cause overfitting. Therefore, selecting features that have majorly contributed to the performance of ML and removing the highly correlated features are necessary to identify relevant features, thus reducing the complexity of models. Among various feature selection methods, we have applied the mRMR (minimum redundancy maximum relevance) method [39], which uses mutual information to compute the relevance and redundancy among features/classes. We have used the mrmr_regression program to select the top 200, 500, 1000, 1500 and 2000 features and evaluated the ML performance on the selected features.

### 2.7. Regression Models

Regression aims to create a model of the relationship between a certain number of features and a continuous target variable. Twelve different regression algorithms were implemented in our study using the Python Scikit-learn library [40]. Regressive algorithms used in the study are Linear regression (LR)[41], Support vector regression (SVR)[42], Ridge regression (Ridge)[43], Lasso regression[44], Gradient Boosting regression (GBR) [45], MLP regression (MLPR) [46], AdaBoost regression(AdaBR) [47], Elastic Net regression (ENR) [48], Kernel Ridge regression (KRR)[49] and Bayesian Ridge regression(BRR)[50] to determine which model could deliver the most accurate performance in the prediction of the exact value of MIC. The models are run with the default parameters.

### 2.8. Hybrid approach

We intended to enhance further our best-performing regressor model, built on the best feature set, by incorporating an ensemble method. This method combines the regressive power of the similarity search method with the ML regressors. Unlike a classification problem where the blast scores are added with the prediction scores, instead here, the MIC values are replaced for the obtained blast hits with the corresponding ML prediction scores. Various E-value cutoffs were used to evaluate the performance of the hybrid method.

### 2.8. Performance metrics

We have used the Correlation coefficient (R-score), Mean absolute percentage error (MAPE), Maximum residual error (MRE), Mean absolute error (MAE), Mean squared error (MSE), Root mean squared error (RMSE), and Coefficient of determination (R²) as metrics to measure the performance of the regressive models as shown in Equations from 2 to 7. Note that the lower the MAE, MSE, MAPE, MRE, or RMSE value, the better the regression models. Conversely, the higher the value of R and R², the better the regressive models.

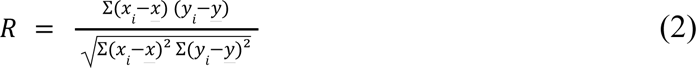

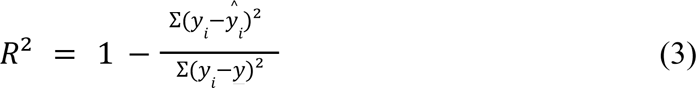

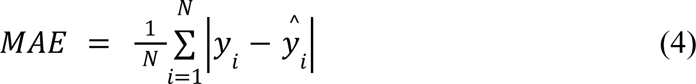

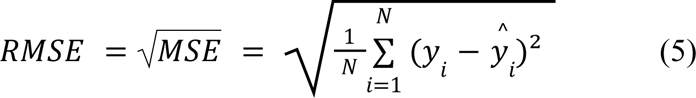

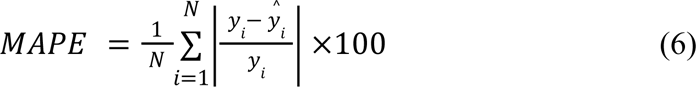

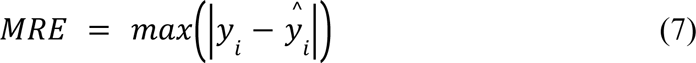

where, *y_i_* and *x_i_* are the data points, N is the number of data points, and <u>*y*</u>, *ŷ_i_* and <u>*x*</u> indicates the mean value of y, predicted value of actual value (y), and average of x, respectively.

## 3. Results

### 3.1. Primary analysis

#### 3.1.1. Length distribution

We have explored the datasets according to their length and found that more than 80% of the data length lies in between 11 to 40 residues for all train, validation and independent sets. The dataset is graphically represented in Figure 3 as per length distribution.

**Figure 3:**
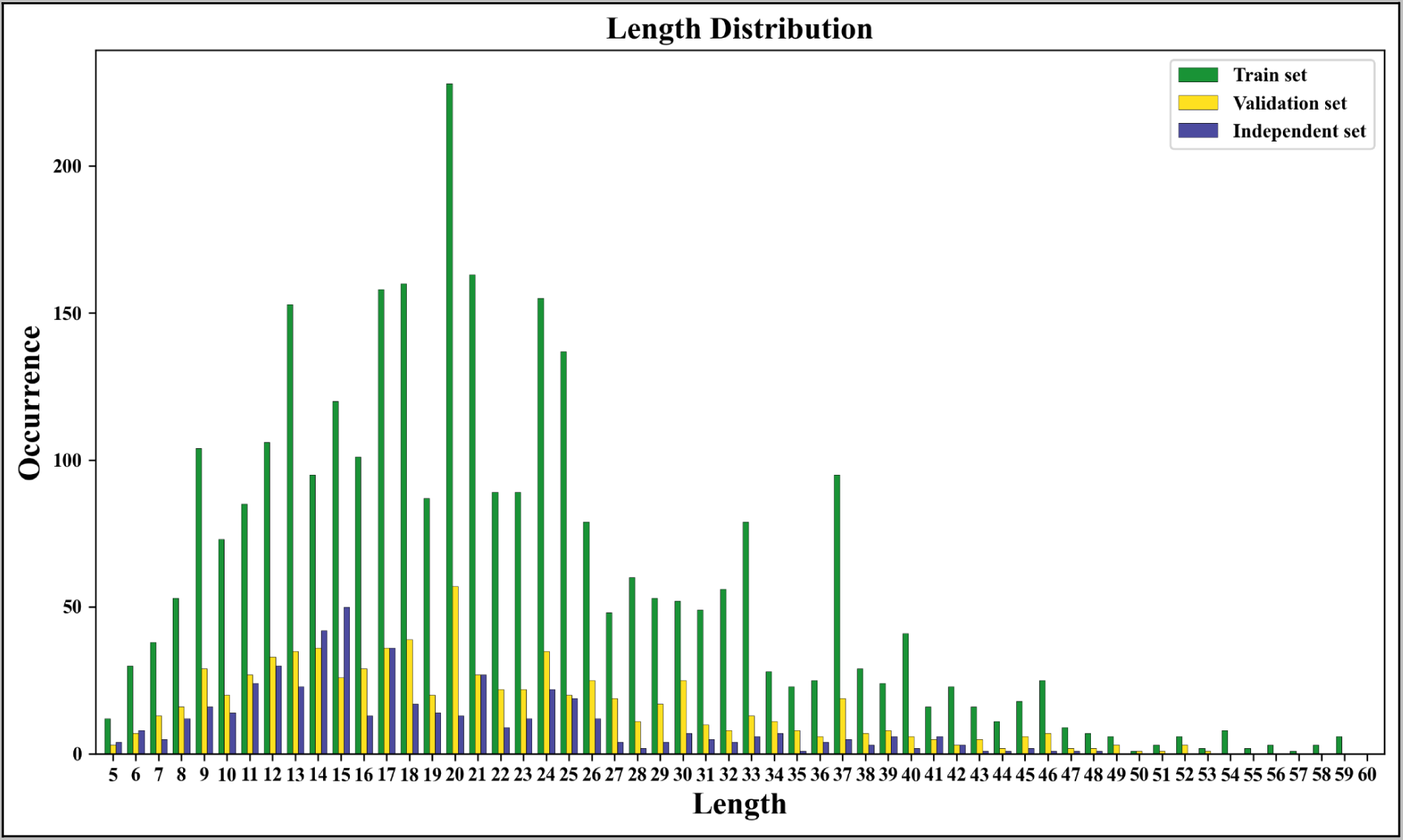
The histogram of the distribution of length of antibacterial peptide for *E. coli* for training, validation and independent set.

#### 3.1.2. Correlation analysis

In an effort to find the strength of the relationship between the features and the target MIC values, we have computed the Pearson correlation of amino acids, dipeptides, and mRMR selected 1000 features, as depicted in Figures 4, 5, and 6. Its value ranges from −1 to 1, where −1 represents a negative correlation, which means the target MIC value is less dependent, and 1 represents a positive correlation, which means the target MIC value is more dependent on that particular feature, and a value of 0 indicates no correlation.

**Figure 4:**
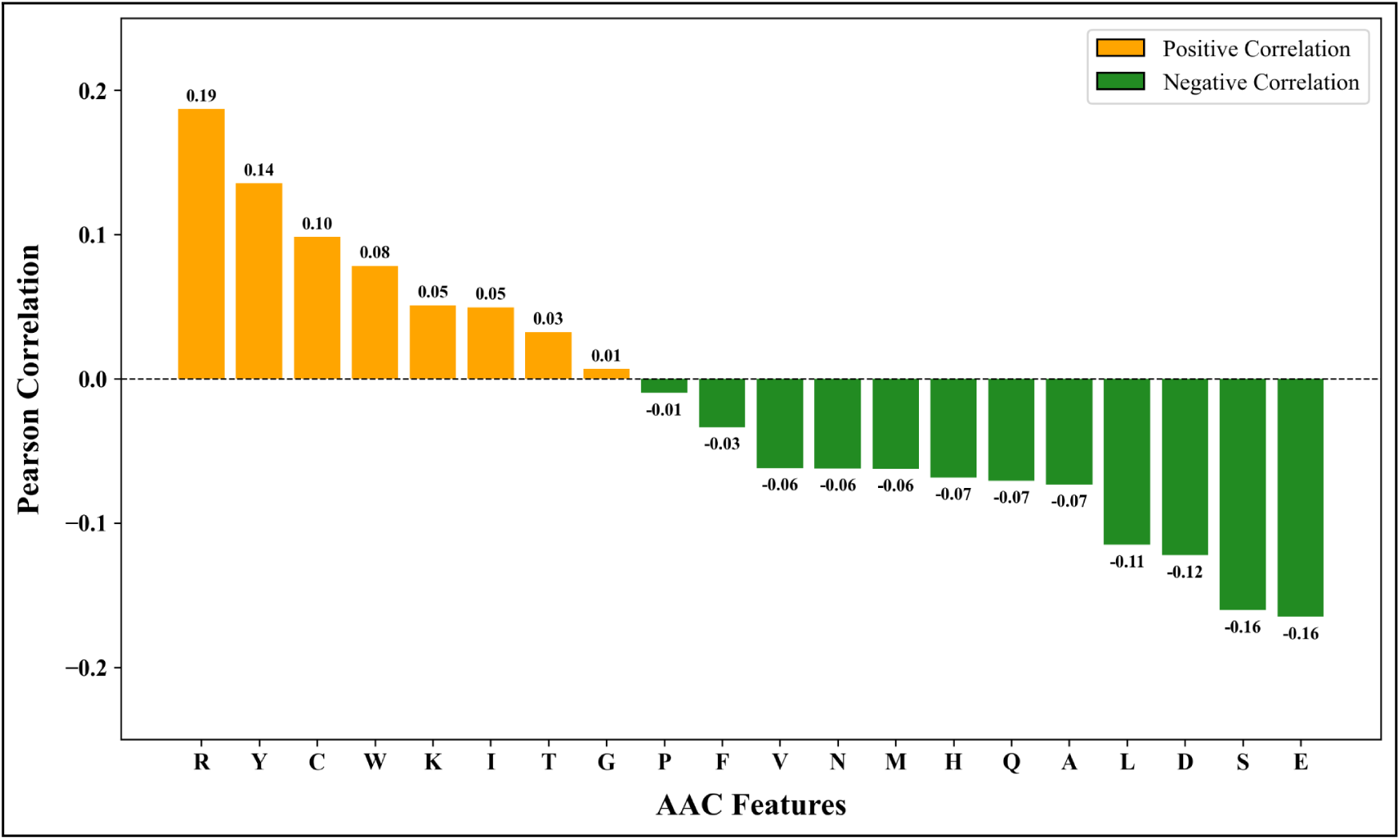
The bar graph showing the Pearson correlation value of AAC features with the target MIC values.

**Figure 5:**
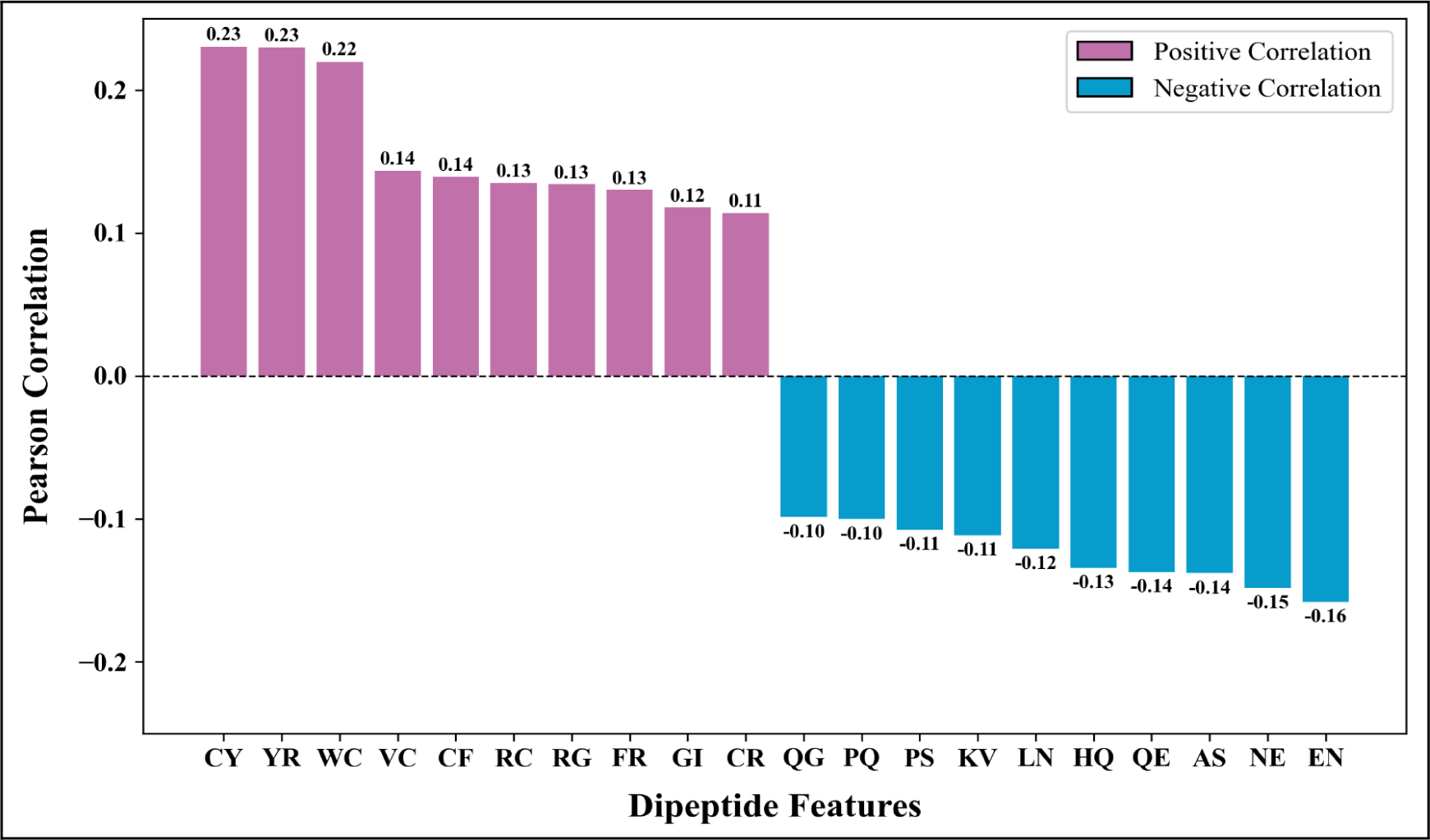
The bar graph showing the Pearson correlation value of Dipeptide features with the target MIC values.

**Figure 6:**
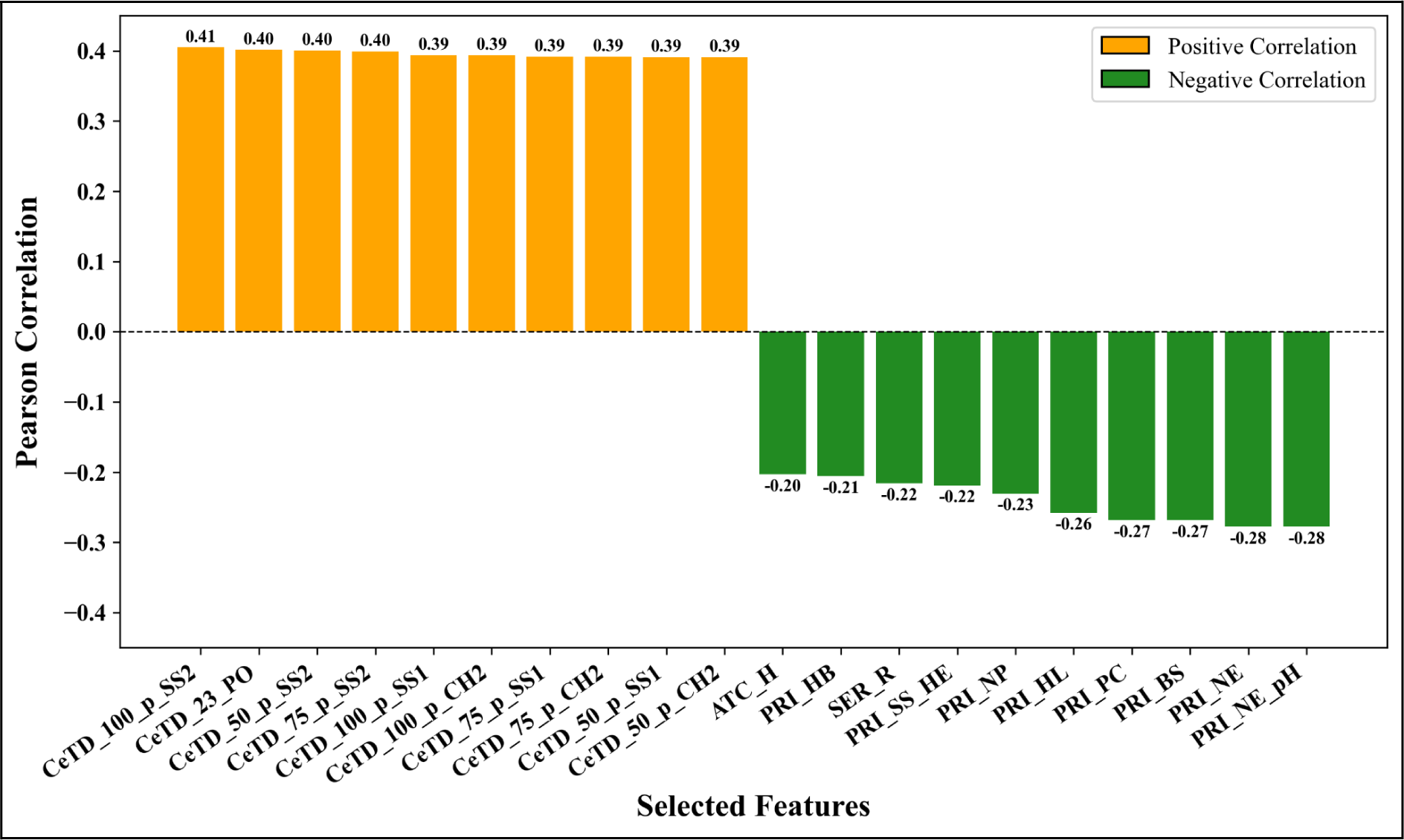
The bar graph showing the Pearson correlation value of 1000 selected features with the target MIC values.

In amino acids, residues such as R and Y show a positive correlation of 0.19 and 0.14, while residues such as E and S show significant negative correlation of −0.16. In the case of dipeptides, among 400 dipeptide features, the top 10 positive correlated and bottom 10 negative correlated features have been selected for representation; dipeptides like CY and YR have shown the highest correlation of 0.23. In contrast, EN and NE have shown the lowest correlation of −0.16 and −0.15, respectively. Similarly, for the 1000 mRMR selected features, the top 10 and bottom 10 have been used for representation, where the CeTD features show a maximum positive correlation of 0.41, and PRI features show a minimum correlation of-0.28.

### 3.2. Performance of Similarity Search Method

We have also utilized the ability of BLAST to predict the MIC values of validation data for the hits obtained from the customized database. For the validation hits obtained, we have assigned them the MIC values corresponding to the database sequences. After that, their performances with respect to the actual MIC values of the validation set were calculated, as shown in Table 2. At the highest e-value, the performance increases, but at the same time, the number of sequence hits obtained by the BLAST is much less, which signifies the inefficiency of the similarity search method in predicting the MIC values of the peptides.

**Table 2:** Performance of BLAST on the validation set at different e-values.

| E-value | Hits<br>(Total peptides =<br>786) | R | R<sup>2</sup> | MSE | RMSE | MAPE | MAE | MRE |
| --- | --- | --- | --- | --- | --- | --- | --- | --- |
| 1.00E-20 | 18 | 0.88 | 0.74 | 0.12 | 0.34 | 1.79 | 0.23 | 0.72 |
| 1.00E-10 | 174 | 0.72 | 0.48 | 0.32 | 0.57 | 15.10 | 0.37 | 2.11 |
| 1.00E-06 | 334 | 0.72 | 0.49 | 0.36 | 0.60 | 104.60 | 0.41 | 2.20 |
| 0.0001 | 406 | 0.70 | 0.45 | 0.37 | 0.60 | 93.80 | 0.41 | 2.84 |
| 0.001 | 461 | 0.69 | 0.41 | 0.39 | 0.62 | 83.53 | 0.42 | 2.84 |
| 0.01 | 516 | 0.70 | 0.43 | 0.38 | 0.61 | 74.72 | 0.41 | 2.84 |
| 0.1 | 570 | 0.69 | 0.42 | 0.39 | 0.63 | 71.75 | 0.42 | 2.84 |
| 1 | 619 | 0.67 | 0.38 | 0.42 | 0.65 | 66.12 | 0.44 | 3.26 |

#R= Correlation coefficient, R<sup>2</sup> = Coefficient of determination, MSE = Mean squared error, RMSE =Root mean squared error, MAPE= Mean absolute percentage error, MAE = Mean absolute error, MRE= Maximum residual error

### 3.3. Performance on ML-based regressors

#### 3.3.1. Compositional features

We have developed prediction models using 12 regression algorithms, including LR, RFR, GBR, ENR, SVR, DTR, Ridge, Lasso, MLPR, AdaBR, KRR, and BRR. We designed prediction models employing all different types of compositional features, as shown in Table 1. Figures 7 and 8 show that the RF-based regressor performs best among all other regression models with the highest R² score and lowest RMSE value using the AAC feature; therefore, we have applied RFR for further analysis. The detailed list of performance of RFR using compositional features is depicted in Table 3; the ALLCOMP feature outperforms other models with an R and R² score of 0.77 and 0.59 on the validation dataset with an RMSE value of 0.53. The additional performance metrics of the ALLCOMP feature are represented in Supplementary Table S2.

**Figure 7:**
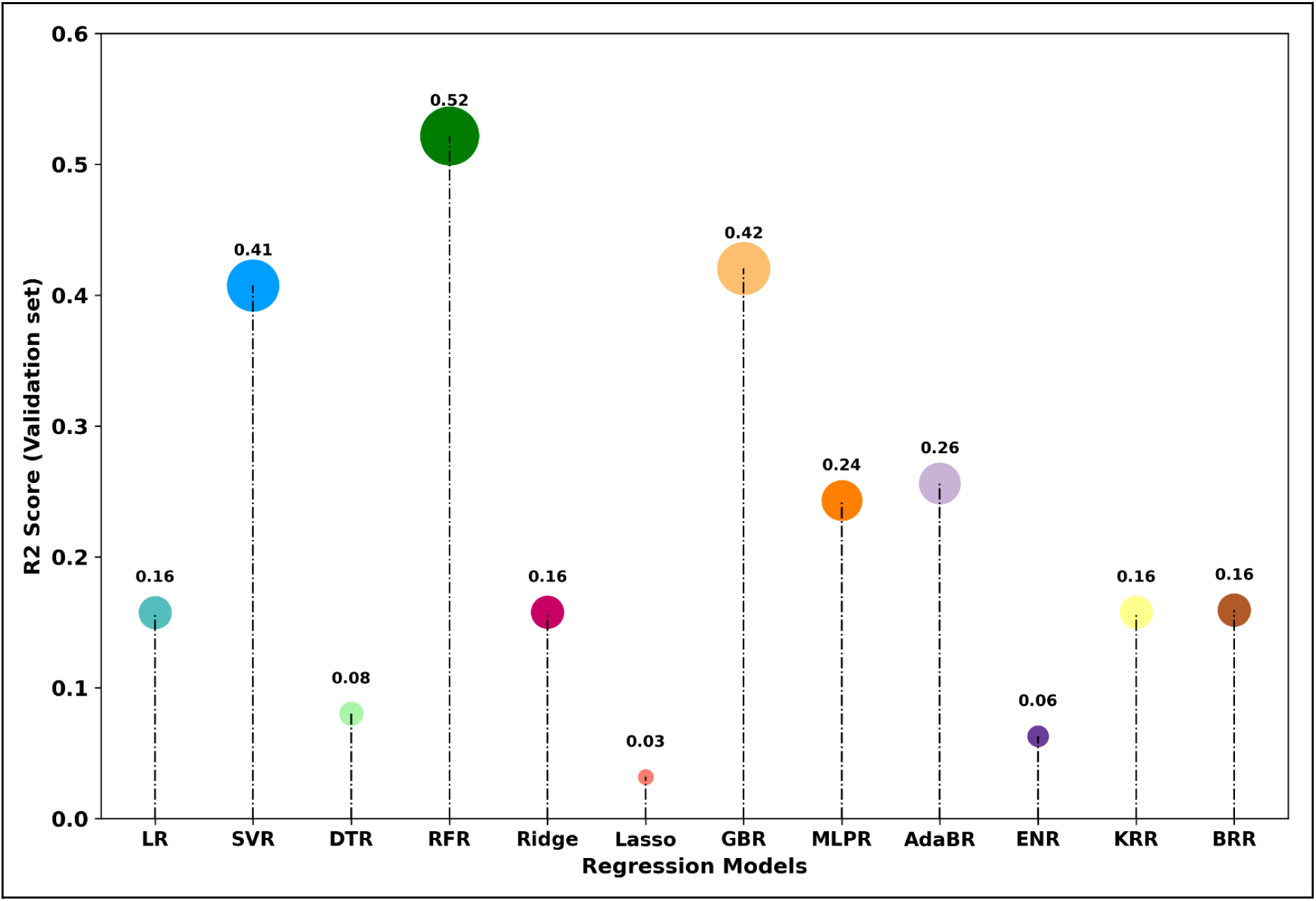
Performance of ML-based regressors in terms of R² scores developed using AAC feature on the validation set using a 5-fold cross-validation strategy.

**Figure 8:**
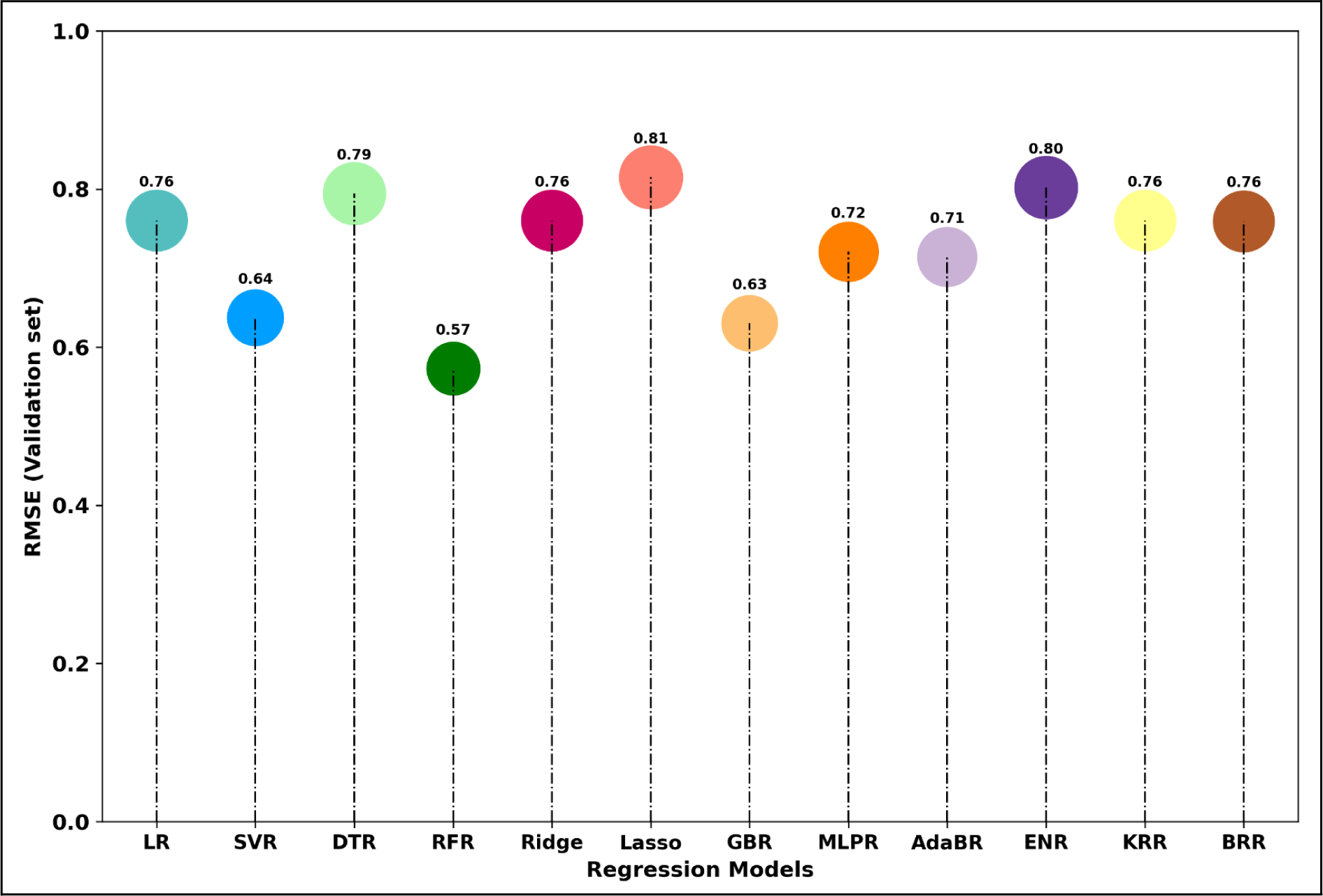
Performance of ML-based regressors in terms of RMSE scores developed using AAC feature on the validation set using a 5-fold cross-validation strategy.

**Table 3:** Performance of RF-based regressor on compositional features of training and validation set.

| Features | R | R<sup>2</sup> | RMSE | R | R<sup>2</sup> | RMSE |
| --- | --- | --- | --- | --- | --- | --- |
|  | TRAINING SET |  |  | VALIDATION SET |  |  |
| AAC | 0.67 | 0.45 | 0.61 | 0.73 | 0.52 | 0.57 |
| AAI | 0.64 | 0.41 | 0.63 | 0.68 | 0.46 | 0.61 |
| APAAC | 0.69 | 0.47 | 0.60 | 0.72 | 0.51 | 0.58 |
| ATC | 0.49 | 0.23 | 0.72 | 0.53 | 0.27 | 0.71 |
| BTC | 0.51 | 0.23 | 0.73 | 0.50 | 0.21 | 0.74 |
| CTC | 0.65 | 0.42 | 0.63 | 0.71 | 0.51 | 0.58 |
| CeTD | 0.70 | 0.49 | 0.59 | 0.76 | 0.57 | 0.55 |
| DDR | 0.66 | 0.44 | 0.62 | 0.71 | 0.49 | 0.59 |
| **DPC** | 0.69 | 0.47 | 0.60 | 0.74 | 0.54 | 0.56 |
| **PAAC** | 0.67 | 0.44 | 0.62 | 0.70 | 0.48 | 0.60 |
| **PCP** | 0.64 | 0.40 | 0.64 | 0.67 | 0.45 | 0.62 |
| **PRI** | 0.66 | 0.43 | 0.62 | 0.70 | 0.48 | 0.60 |
| **QSO** | 0.68 | 0.47 | 0.60 | 0.74 | 0.54 | 0.56 |
| **RRI** | 0.63 | 0.40 | 0.64 | 0.67 | 0.45 | 0.62 |
| **SEP** | 0.27 | -0.07 | 0.85 | 0.30 | -0.03 | 0.84 |
| **SER** | 0.68 | 0.46 | 0.61 | 0.73 | 0.52 | 0.57 |
| **SOC** | 0.28 | 0.00 | 0.82 | 0.32 | 0.04 | 0.81 |
| **SPC** | 0.63 | 0.40 | 0.64 | 0.67 | 0.44 | 0.62 |
| **TPC** | 0.67 | 0.45 | 0.61 | 0.72 | 0.52 | 0.57 |
| **ALLCOMP** | **0.72** | **0.52** | **0.57** | **0.77** | **0.59** | **0.53** |

#R= Correlation coefficient, R<sup>2</sup> = Coefficient of determination, RMSE =Root mean squared error

#### 3.3.2. Binary Profiles

To understand the importance of building different models for different length sets, we have developed binary profiles for different length groups and used them as input features to train and evaluate the prediction models by implementing various ML regressors. As shown in Table 4, the RFR performs extremely poor in the length group of 41 or more residues with the highest RMSE value on the validation set, while it performs better for 11-20 residues with an R² and RMSE value of 0.55 and 0.57 on the validation set. The detailed results of binary profiles for all length groups are depicted in Supplementary Tables S3, S4, S5, S6 and S7.

**Table 4:** Performance of RF-based regressor on Binary profiles of different length groups of training and validation set.

| Length groups | R | R<sup>2</sup> | RMSE | R | R<sup>2</sup> | RMSE |
| --- | --- | --- | --- | --- | --- | --- |
|  | TRAINING SET |  |  | VALIDATION SET |  |  |
| **5-10aa** | 0.67 | 0.44 | 0.69 | 0.73 | 0.53 | 0.63 |
| **11-20aa** | 0.70 | 0.49 | 0.6 | 0.74 | 0.55 | 0.57 |
| **21-40aa** | 0.60 | 0.36 | 0.58 | 0.53 | 0.28 | 0.6 |
| **41 or more aa** | 0.55 | 0.21 | 0.67 | 0.48 | 0.21 | 0.69 |
| **Whole data** | 0.59 | 0.35 | 0.66 | 0.64 | 0.41 | 0.64 |

#R= Correlation coefficient, R<sup>2</sup> = Coefficient of determination, RMSE =Root mean squared error

#### 3.3.3. LLM embeddings

To generate the embedding using large language models, first, we have to modify the LL models for regression analysis and then pass out data to compute the embeddings. The performance across various LLM embeddings is reported in Table 5. Models that exploited ESM-2 embeddings saw the best performance with lower RMSE and high correlation scores of 0.56 and 0.76, respectively. Supplementary Tables S8, S9, and S10 show the results obtained with all three LLM embeddings.

**Table 5:** Performance of RF-based regressor on LLM embeddings of training and validation set.

| LLM | R | R<sup>2</sup> | RMSE | R | R<sup>2</sup> | RMSE |
| --- | --- | --- | --- | --- | --- | --- |
|  | TRAINING SET |  |  | VALIDATION SET |  |  |
| **ProtBERT** | 0.65 | 0.40 | 0.64 | 0.70 | 0.46 | 0.61 |
| **BioBERT** | 0.58 | 0.31 | 0.69 | 0.63 | 0.36 | 0.66 |
| **ESM-2** | 0.69 | 0.46 | 0.61 | 0.76 | 0.55 | 0.56 |

#R= Correlation coefficient, R<sup>2</sup> = Coefficient of determination, RMSE =Root mean squared error

### 3.4. Performance on selected features

We have implemented the mRMR feature selection technique to get useful features that help enhance the models’ performances. We have selected the top 200, 500, 1000,1500 and 2000 features to check the effectiveness of different feature subsets. The performance of the RF-based regressor on selected features is depicted in Table 6, which shows that the top 1000 features contributed most to effective understanding by the model and got a maximum R/R^2^of 0.78/0.59 with RMSE 0.53 on the validation set. The detailed performance of the selected feature subsets are presented in Supplementary Tables S11, S12, S13, S14 and S15.

**Table 6:**
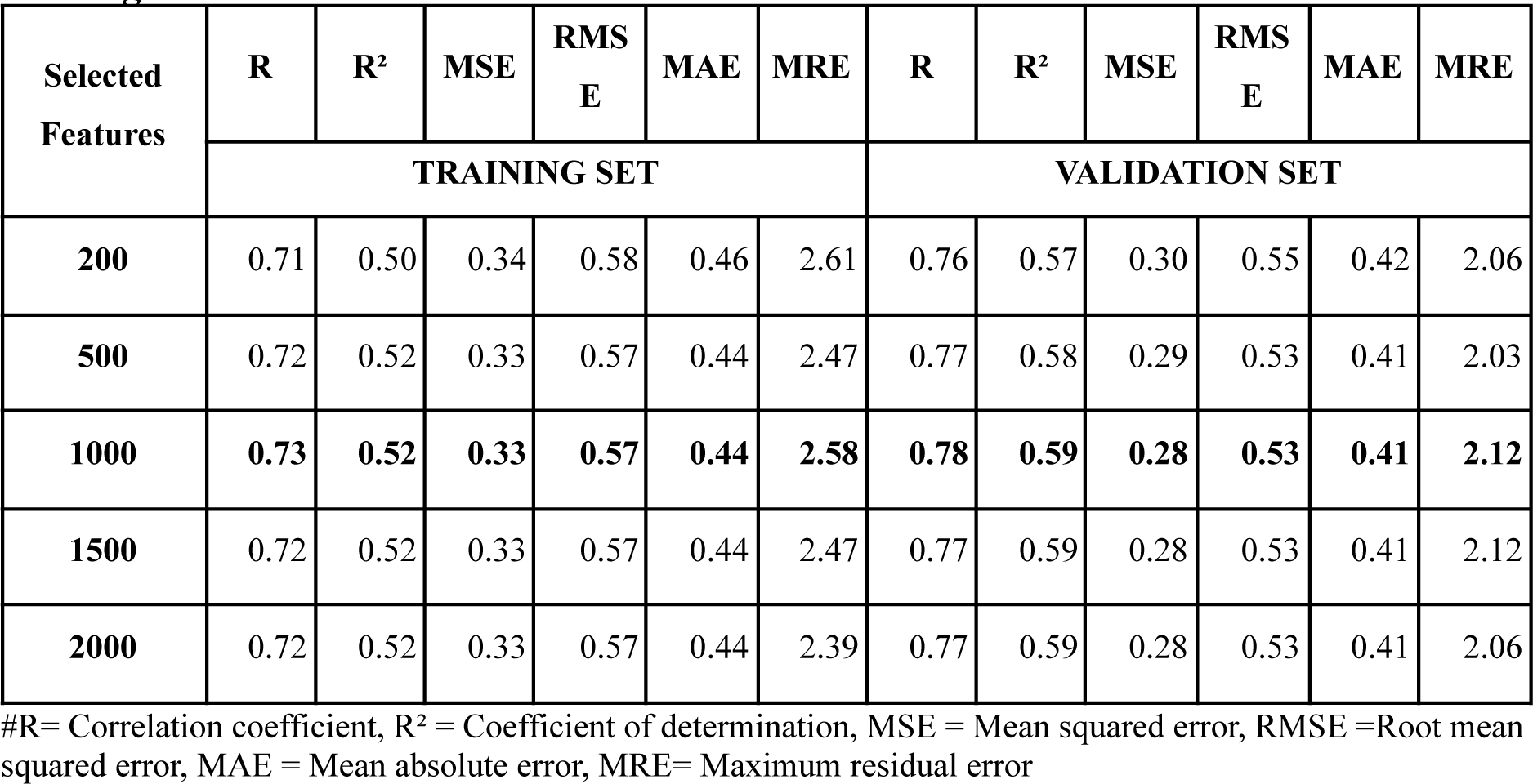
Performance of RF-based regressor on different selected feature subsets on training and validation set.

| Selected Features | R | R<sup>2</sup> | MSE | RMS E | MAE | MRE | R | R<sup>2</sup> | MSE | RMS E | MAE | MRE |
| --- | --- | --- | --- | --- | --- | --- | --- | --- | --- | --- | --- | --- |
|  | TRAINING SET |  |  |  |  |  | VALIDATION SET |  |  |  |  |  |
| 200 | 0.71 | 0.50 | 0.34 | 0.58 | 0.46 | 2.61 | 0.76 | 0.57 | 0.30 | 0.55 | 0.42 | 2.06 |
| 500 | 0.72 | 0.52 | 0.33 | 0.57 | 0.44 | 2.47 | 0.77 | 0.58 | 0.29 | 0.53 | 0.41 | 2.03 |
| 1000 | 0.73 | 0.52 | 0.33 | 0.57 | 0.44 | 2.58 | 0.78 | 0.59 | 0.28 | 0.53 | 0.41 | 2.12 |
| 1500 | 0.72 | 0.52 | 0.33 | 0.57 | 0.44 | 2.47 | 0.77 | 0.59 | 0.28 | 0.53 | 0.41 | 2.12 |
| 2000 | 0.72 | 0.52 | 0.33 | 0.57 | 0.44 | 2.39 | 0.77 | 0.59 | 0.28 | 0.53 | 0.41 | 2.06 |

#R= Correlation coefficient, R<sup>2</sup> = Coefficient of determination, MSE = Mean squared error, RMSE =Root mean squared error, MAE = Mean absolute error, MRE= Maximum residual error

### 3.5. Performance of Hybrid Method

With the aim of improving the performance of the ML-based regressors, we have tried a hybrid approach that combines the similarity search BLAST with the non-similarity-based ML methods. Here, we first run the BLAST on the validation set and obtain the hits from the customized database. After that, for the validation hits obtained, we assigned them the MIC values as predicted by the best-performing RF-based regressor model built on mRMR-selected 1000 features and their performances with respect to the actual MIC values of the validation set were calculated, as shown in Table 7. Our RFR model obtained an R score of 0.78 on the validation set, while the hybrid method only for obtained hits shows a maximum R score of 0.92 at an e-value of 1.00E-20 while its coverage is only 18 peptides from the total 786 peptides, which is very low implies that the combined approach is not improving the performance significantly.

**Table 7:** Performance of hybrid method on the validation set at different e-values.

| **E-value** | **Hits**<br>**(Total peptides =**<br>**786)** | **R** | **R<sup>2</sup>** | **MSE** | **RMSE** | **MAPE** | **MAE** | **MRE** |
| --- | --- | --- | --- | --- | --- | --- | --- | --- |
| **1.00E-20** | 18 | 0.92 | 0.76 | 0.11 | 0.32 | 3.05 | 0.27 | 0.68 |
| **1.00E-10** | 174 | 0.76 | 0.54 | 0.29 | 0.54 | 34.12 | 0.40 | 2.12 |
| **1.00E-06** | 334 | 0.80 | 0.61 | 0.27 | 0.52 | 200.39 | 0.40 | 2.12 |
| **0.0001** | 406 | 0.78 | 0.59 | 0.27 | 0.52 | 168.97 | 0.40 | 2.12 |
| **0.001** | 461 | 0.77 | 0.58 | 0.28 | 0.53 | 149.32 | 0.41 | 2.12 |
| **0.01** | 516 | 0.78 | 0.59 | 0.27 | 0.52 | 133.57 | 0.40 | 2.12 |
| **0.1** | 570 | 0.78 | 0.60 | 0.27 | 0.52 | 122.52 | 0.41 | 2.12 |
| **1** | 619 | 0.78 | 0.60 | 0.28 | 0.53 | 112.86 | 0.41 | 2.12 |
| **ML (RFR-1000 Features)** |  | **0.78** | **0.59** | **0.28** | **0.53** | **89.25** | **0.41** | **2.12** |

#R= Correlation coefficient, $R^2$ = Coefficient of determination, MSE = Mean squared error, RMSE = Root mean squared error, MAPE = Mean absolute percentage error, MAE = Mean absolute error, MRE= Maximum residual error, ML = Machine learning, RFR = Random Forest Regression

### 3.6. Benchmarking on an independent set

We have selected an RF-based regressor-based model on default parameters with the top 1000 selected features as our final model. To evaluate the performance of our model, we have utilized the independent set and compared our performance with the existing methods. The MBC-attention model[23] was built using CNN with an attention layer and tuned on various hyperparameters such as the number of layers, number of filters, dropout layers, loss function and early stopping parameters. We have used the same non-redundant dataset that is used in MBC-attention. However, the dataset splitting is performed differently. Similarly, the model architecture of AMPActiPred is the deep neural network that first classifies the ABP or non-ABP against different bacterial groups and finally predicts the activity of those peptides in terms of MIC values [22]. The performance of different models on the independent dataset is shown in Table 8, which suggests that our model performs better on the independent set despite being built using basic computational procedures with an R² and RMSE value of 0.410 and 0.709, respectively.

**Table 8:** Comparative performance of our models and existing methods on the independent dataset.

| Model | R | $R^2$ | MSE | RMSE | MAPE | MAE | MRE |
| --- | --- | --- | --- | --- | --- | --- | --- |
| **Our models** |  |  |  |  |  |  |  |
| **EIPpred (1000 selected features-RFR)** | **0.676** | **0.41** | **0.503** | **0.709** | **5.975** | **0.567** | **2.362** |
| **ALLCOMP (RFR)** | 0.673 | 0.401 | 0.511 | 0.715 | 5.968 | 0.572 | 2.335 |
| **ESM-2 (RFR)** | 0.611 | 0.326 | 0.575 | 0.758 | 6.571 | 0.591 | 2.944 |
| **Existing models** |  |  |  |  |  |  |  |
| **MBC-attention (CNN)** | 0.657 | 0.392 | 0.519 | 0.72 | 7.436 | 0.554 | 2.763 |
| **AMPActiPred (DNN)** | 0.334 | -0.071 | 0.914 | 0.956 | 6.298 | 0.585 | 4.258 |

#R= Correlation coefficient, R<sup>2</sup> = Coefficient of determination, MSE = Mean squared error, RMSE =Root mean squared error, MAPE= Mean absolute percentage error, MAE = Mean absolute error, MRE= Maximum residual error, RFR = Random Forest Regression, CNN = Convolutional Neural Network, DNN = Deep Neural Network

### 3.7. Web server implementation

To better serve the scientific community, we have developed a user-friendly prediction web interface named “EIPpred” (https://webs.iiitd.edu.in/raghava/eippred) and executed our best model to predict the MIC values of ABPs. The web server includes three different modules: “Predict”, “Design”, and “Protein scan”. The module ‘Predict’ allows users to predict the inhibitory activity of the submitted sequence against *E. coli* in terms of MIC value (in −log₁₀ µM). The ‘Design’ module allows users to create mutants by defining the position of mutation and residue to be mutated in the input sequence. The result of the design module includes the mutant sequences with inhibitory activity and represents their predicted MIC values along with the correlation plot for the original and mutant sequence in order to better understand the importance of each residue in imparting the inhibitory activity to the peptide. The ‘Protein scan’ module scans the protein sequences to identify the inhibitory regions/domains based on the predicted MIC values against *E. coli* of the desired lengths defined by the user. We also allow users to download the training, validation, and independent dataset (https://webs.iiitd.edu.in/raghava/eippred/dataset.php), which we have used in this study, available in CSV file format.

## 4. Discussion

Bacteria are well-studied organisms in terms of their structure, biology, and infection processes; nevertheless, our understanding remains incomplete due to their fast progression. The rat race between bacterial evolution and antibiotic research demands alternative options to overcome resistance that directly targets bacterial survival pathways, such as peptide-based therapies. *E. coli*, a common clinical infection, frequently develops resistance to several antimicrobials, creating an enormous global issue [51]. Thus, predicting the inhibitory activity of peptides, specifically through MIC values, is crucial for assessing drug effectiveness against specific bacterial species.

Since existing approaches are computationally expensive and have limited species-specificity, the activity prediction of ABPs requires improvement; hence, designing newer inhibitory peptides has become essential in the post-antibiotic era. In this study, we report a quick and efficient activity RF-based regressor tool for ABPs to improve the prediction of quantitative MIC values of ABPs against *E. coli*. Our model relies on a supervised ML approach based on sequence information employing various types of feature sets. The main advantage of our model is its simple architecture with superior performance that allows us to learn from multiple feature types. This is particularly useful in the case where employing heavy computation is not possible.

The initial analysis of the correlation of the selected 1000 features with the target MIC values shows a strong correlation with the CeTD feature, implying that the distribution of group 1 & 2 residues for the secondary structure (SS), charge(CH) and transition of group 2 to 3 residues for polarity (PO) along the whole length of the sequence are important in defining the inhibitory properties of the ABPs (The residues in SS group 1 are E, A, L, M, Q, K, R, and H and in group 2 are V, I, Y, C, W, F, and T; the residues in CH group 1 are K, R and group 2 are A, N, C, Q, G, H, I, L, M, F, P, S, T, W, Y, V & the residues in PO group 2 are P, A, T, G, S and group 3 are H, Q, R, K, N, E, D). The residue repeats for physio-chemical properties, such as neutral, basic and positively charged residues, negatively correlate with the target variable. This implies that the distribution of residues capable of forming secondary structures is essential in conferring inhibitory activity.

In this study, we leverage the regressive ability of various ML algorithms for quantitative MIC prediction and enhance their predictive capabilities by incorporating different feature sets such as compositional, binary profiles, and LLM embedding. Among the features set used, the model built using all compositional features performs significantly better than individual compositional features with an R and R² score of 0.77 and 0.59, respectively, on the validation set. Secondly, the binary profiles of different length groups suggest that the regressive models built using peptides with lengths from 5 to 20 residues are better at capturing the properties of the peptide to predict the MIC values quantitatively. Third and lastly, ESM-2 performs well but not better than all compositional feature sets among the LLM embedding generated from the pre-trained models.

Since the number of all compositional features is high, we employed mRMR to select the features that greatly contributed to the performance of the model and from Table 6, we conclude that the top 1000 feature set is performing superior to other selected feature sets, thus reducing overfitting and irrelevant features. Moreover, we also perform BLAST to understand the effect of the similarity on predicting the activity of the peptides; however, our results suggest that utilizing BLAST hits doesn’t significantly improve the performance of our models and also the prediction given by our ML models for the hits is far better than validation hits MIC values replaced by the subject MICs as done in the hybrid method shown in Table 7.

It is important to note that by performing the comparative analysis on the independent dataset, our final RFR model built using 1000 selected features outperforms the existing methods by only small margins, achieving R and RMSE values of 0.676 and 0.709, still employing simple architecture instead of sophisticated neural networks, greatly enhancing the performance of our model. This reduces the cost and time needed to predict the activity of ABPs in *E. coli.* Lastly, we built a web server - EIPpred and in addition to the web server facility, GitHub (https://github.com/NishaBajiya/EIPpred), the standalone version, and the pip package (https://pypi.org/project/eippred/) of our model are also publicly available to researchers. By utilizing our web server, novel inhibitory ABPs can be designed against *E. coli* as defined by the user with their predicted MIC values. Eventually, the model needs more improvement and extends its domain to predict the activity of peptides for other pathogenic species.

## Authors’ contributions

NB collected the dataset. NB and GPSR processed the datasets. NB implemented the algorithms and developed the prediction models. NB and GPSR analyzed the results. NB created the front-end user interface, and NK created the back-end of the web server and standalone package. NB and GPSR performed the writing, reviewing, and draft preparation of the manuscript. GPSR conceived and coordinated the project and gave overall supervision to the project. All authors have read and approved the final manuscript.

## Acknowledgements

Authors are thankful to the Council of Scientific & Industrial Research (CSIR), University Grants Commission (UGC) and Department of Bio-Technology (DBT) for fellowships and financial support, and the Department of Computational Biology, IIITD New Delhi for infrastructure and facilities. We would like to acknowledge that the Figures were created using BioRender.

## Data Availability

The dataset (https://webs.iiitd.edu.in/raghava/eippred/dataset.php) and code files (https://webs.iiitd.edu.in/raghava/eippred/standalone.php) used in EIPpred are available on the web server.

## Author’s Biography

1. Nisha Bajiya is currently working as Ph.D. in Computational Biology from Department of Computational Biology, Indraprastha Institute of Information Technology, New Delhi, India.
2. Nishant Kumar is currently working as Ph.D. in Computational biology from Department of Computational Biology, Indraprastha Institute of Information Technology, New Delhi, India.
3. Gajendra P. S. Raghava is currently working as Professor and Head of Department of Computational Biology, Indraprastha Institute of Information Technology, New Delhi, India.

## Abbreviation

ABP: Antibacterial peptide
R: Correlation coefficient
R^2^: Coefficient of determination
MSE: Mean squared error
RMSE: Root mean squared error
MAPE: Mean absolute percentage error
MAE: Mean absolute error
MRE: Maximum residual error
BLAST: Basic Local Alignment Search Tool
mRMR: Minimum Redundancy Maximum Relevance
RFR: Random Forest Regression
CNN: Convolutional Neural Network
DNN: Deep Neural Network

## Supplementary Legends

**Supplementary Table S1: Summary of distribution of sequence length groups for Binary profiles**

**Supplementary Table S2: Performance of ALLCOMP feature of training and validation set on different regression models**

**Supplementary Table S3: Performance of Binary profile-based regressors for the whole length of training and validation set**

**Supplementary Table S4: Performance of Binary profile-based regressors for length group 5-10 residues on training and validation set**

**Supplementary Table S5: Performance of Binary profile-based regressors for length group 11-20 residues on training and validation set**

**Supplementary Table S6: Performance of Binary profile-based regressors for length group 21-40 residues on training and validation set**

**Supplementary Table S7: Performance of Binary profile-based regressors for length group 41 or more residues on training and validation set**

**Supplementary Table S8: Performance of ProtBERT embeddings-based regressors on training and validation set**

**Supplementary Table S9: Performance of BioBERT embeddings-based regressors on training and validation set**

**Supplementary Table S10: Performance of ESM-2 embeddings-based regressors on training and validation set**

**Supplementary Table S11: Performance of RF-based regressor on Top 200 selected features on training and validation set**

**Supplementary Table S12: Performance of RF-based regressor on Top 500 selected features on training and validation set**

**Supplementary Table S13: Performance of RF-based regressor on Top 1000 selected features on training and validation set**

**Supplementary Table S14: Performance of RF-based regressor on Top 1500 selected features on training and validation set**

**Supplementary Table S15: Performance of RF-based regressor on Top 2000 selected features on training and validation set**

