## Supplementary Table for "Prediction of inhibitory peptides against *E. coli* with desired MIC value"

**Mailing Address of Authors**

***Corresponding Author**

Prof. Gajendra P. S. Raghava

Head and Professor

Department of Computational Biology

Indraprastha Institute of Information Technology, Delhi

Okhla Industrial Estate, Phase III (Near Govind Puri Metro Station)

New Delhi, India – 110020 Office: A-302 (R&D Block)

Website: <http://webs.iiitd.edu.in/raghava/>

**Supplementary Table S1: Summary of distribution of sequence length groups for Binary profiles**

| **Length groups** | **Fixed length vector** | **Binary profiles** | **Total sequences** | **Training set** | **Validation set** | **Independent set** |
| --- | --- | --- | --- | --- | --- | --- |
| **5-10aa** | 10 | 10 * 20 | 398 | 318 | 80 | 59 |
| **11-20aa** | 20 | 20 * 20 | 1631 | 1304 | 327 | 262 |
| **21-40aa** | 40 | 40 * 20 | 1693 | 1354 | 339 | 159 |
| **41or more aa** | 80 | 80 * 20 | 207 | 165 | 42 | 18 |
| **Whole data** | 10 | 10 * 20 | 3929 | 3143 | 786 | 498 |

**Supplementary Table S2: Performance of ALLCOMP feature of training and validation set on different regression models**

| **Model** | **R** | **R2** | **MSE** | **RMSE** | **MAPE** | **MAE** | **MRE** | **R** | **R2** | **MSE** | **RMSE** | **MAPE** | **MAE** | **MRE** |
| --- | --- | --- | --- | --- | --- | --- | --- | --- | --- | --- | --- | --- | --- | --- |
|  | **TRAINING SET** | | | | | | | **VALIDATION SET** | | | | | | |
| **LR** | 0.02 | -38713.47 | 27073.21 | 134.52 | 3867.75 | 88.46 | 981.86 | 0.01 | -105478.03 | 72365.79 | 269.01 | 29911.13 | 184.92 | 1383.13 |
| **SVR** | 0.42 | 0.16 | 0.57 | 0.76 | 134.95 | 0.61 | 2.67 | 0.40 | 0.16 | 0.58 | 0.76 | 73.38 | 0.61 | 3.06 |
| **DTR** | 0.52 | 0.04 | 0.65 | 0.81 | 107.31 | 0.59 | 3.58 | 0.57 | 0.14 | 0.59 | 0.77 | 112.94 | 0.56 | 4.20 |
| **RFR** | **0.72** | **0.52** | **0.33** | **0.57** | **77.97** | **0.44** | **2.34** | **0.77** | **0.59** | **0.28** | **0.53** | **90.55** | **0.41** | **2.08** |
| **Ridge** | 0.43 | -1.02 | 1.38 | 1.17 | 53.50 | 0.85 | 6.59 | 0.37 | -1.11 | 1.45 | 1.20 | 501.06 | 0.88 | 5.34 |
| **Lasso** | 0.42 | 0.17 | 0.56 | 0.75 | 115.80 | 0.61 | 2.66 | 0.49 | 0.23 | 0.53 | 0.73 | 127.80 | 0.58 | 2.84 |
| **GBR** | 0.67 | 0.44 | 0.38 | 0.62 | 75.92 | 0.49 | 2.34 | 0.72 | 0.51 | 0.34 | 0.58 | 111.22 | 0.46 | 2.20 |
| **MLPR** | 0.52 | -0.08 | 0.74 | 0.86 | 71.93 | 0.64 | 4.62 | 0.47 | -0.17 | 0.81 | 0.90 | 256.45 | 0.66 | 4.67 |
| **AdaBR** | 0.59 | 0.32 | 0.46 | 0.68 | 101.28 | 0.56 | 2.31 | 0.63 | 0.37 | 0.43 | 0.66 | 114.36 | 0.54 | 2.11 |
| **ENR** | 0.46 | 0.20 | 0.54 | 0.74 | 111.89 | 0.59 | 2.68 | 0.53 | 0.27 | 0.50 | 0.71 | 113.54 | 0.56 | 2.79 |
| **KRR** | 0.43 | -1.02 | 1.38 | 1.17 | 53.55 | 0.85 | 6.59 | 0.37 | -1.11 | 1.45 | 1.20 | 501.05 | 0.88 | 5.34 |
| **BRR** | 0.67 | 0.44 | 0.38 | 0.62 | 53.73 | 0.48 | 2.50 | 0.70 | 0.49 | 0.35 | 0.59 | 127.86 | 0.46 | 2.34 |

#R = Correlation coefficient, R² = Coefficient of determination, MSE = Mean squared error, RMSE = Root mean squared error, MAPE = Mean absolute percentage error, MAE = Mean absolute error, MRE = Maximum residual error, LR = Linear regression, SVR = Support vector regression, Ridge = Ridge regression, Lasso = =Lasso regression, GBR = Gradient Boosting regression, MLPR = MLP regression , AdaBR = AdaBoost regression, ENR = Elastic Net regression, KRR = Kernel Ridge regression and BRR = Bayesian Ridge regression

**Supplementary Table S3: Performance of Binary profile-based regressors for the whole length of training and validation set**

| **Model** | **R** | **R2** | **MSE** | **RMSE** | **MAPE** | **MAE** | **MRE** | **R** | **R2** | **MSE** | **RMSE** | **MAPE** | **MAE** | **MRE** |
| --- | --- | --- | --- | --- | --- | --- | --- | --- | --- | --- | --- | --- | --- | --- |
|  | **TRAINING SET** | | | | | | | **VALIDATION SET** | | | | | | |
| **LR** | 0.45 | 0.18 | 0.56 | 0.75 | 86.14 | 0.59 | 2.57 | 0.49 | 0.23 | 0.53 | 0.73 | 112.78 | 0.58 | 2.55 |
| **SVR** | 0.61 | 0.37 | 0.43 | 0.66 | 98.77 | 0.51 | 2.52 | 0.68 | 0.45 | 0.37 | 0.61 | 110.40 | 0.48 | 2.21 |
| **DTR** | 0.42 | -0.14 | 0.77 | 0.88 | 74.07 | 0.64 | 3.32 | 0.44 | -0.08 | 0.74 | 0.86 | 220.66 | 0.62 | 4.16 |
| **RFR** | 0.59 | 0.35 | 0.44 | 0.66 | 75.31 | 0.52 | 2.54 | 0.64 | 0.41 | 0.41 | 0.64 | 122.03 | 0.50 | 2.18 |
| **Ridge** | 0.46 | 0.20 | 0.55 | 0.74 | 95.53 | 0.59 | 2.57 | 0.49 | 0.24 | 0.52 | 0.72 | 115.55 | 0.57 | 2.54 |
| **Lasso** | 0.00 | -0.01 | 0.69 | 0.83 | 131.08 | 0.67 | 2.87 | NaN | 0.00 | 0.69 | 0.83 | 172.03 | 0.67 | 2.96 |
| **GBR** | 0.52 | 0.26 | 0.51 | 0.71 | 106.46 | 0.57 | 2.51 | 0.57 | 0.30 | 0.48 | 0.69 | 140.16 | 0.56 | 2.56 |
| **MLPR** | 0.55 | 0.24 | 0.52 | 0.72 | 90.76 | 0.55 | 2.84 | 0.63 | 0.35 | 0.44 | 0.67 | 189.16 | 0.52 | 2.41 |
| **AdaBR** | 0.35 | 0.09 | 0.62 | 0.79 | 120.39 | 0.64 | 2.63 | 0.39 | 0.11 | 0.61 | 0.78 | 157.59 | 0.63 | 2.91 |
| **ENR** | 0.00 | -0.01 | 0.69 | 0.83 | 131.08 | 0.67 | 2.87 | NaN | 0.00 | 0.69 | 0.83 | 172.03 | 0.67 | 2.96 |
| **KRR** | 0.46 | 0.20 | 0.55 | 0.74 | 95.17 | 0.59 | 2.57 | 0.49 | 0.24 | 0.52 | 0.72 | 115.57 | 0.57 | 2.54 |
| **BRR** | 0.47 | 0.22 | 0.53 | 0.73 | 98.42 | 0.58 | 2.56 | 0.51 | 0.26 | 0.51 | 0.71 | 119.63 | 0.57 | 2.51 |

#R = Correlation coefficient, R² = Coefficient of determination, MSE = Mean squared error, RMSE = Root mean squared error, MAPE = Mean absolute percentage error, MAE = Mean absolute error, MRE = Maximum residual error, LR = Linear regression, SVR = Support vector regression, Ridge = Ridge regression, Lasso = =Lasso regression, GBR = Gradient Boosting regression, MLPR = MLP regression , AdaBR = AdaBoost regression, ENR = Elastic Net regression, KRR = Kernel Ridge regression and BRR = Bayesian Ridge regression

**Supplementary Table S4: Performance of Binary profile-based regressors for length group 5-10 residues on training and validation set**

| **Model** | **R** | **R2** | **MSE** | **RMSE** | **MAPE** | **MAE** | **MRE** | **R** | **R2** | **MSE** | **RMSE** | **MAPE** | **MAE** | **MRE** |
| --- | --- | --- | --- | --- | --- | --- | --- | --- | --- | --- | --- | --- | --- | --- |
|  | **TRAINING SET** | | | | | | | **VALIDATION SET** | | | | | | |
| **LR** | 0.40 | -2.03 | 2.60 | 1.61 | 1.42 | 1.15 | 5.44 | 0.34 | -1.99 | 2.55 | 1.60 | 1.13 | 1.07 | 5.67 |
| **SVR** | 0.62 | 0.39 | 0.53 | 0.73 | 0.94 | 0.57 | 1.94 | 0.70 | 0.48 | 0.44 | 0.66 | 0.82 | 0.52 | 1.65 |
| **DTR** | 0.65 | 0.35 | 0.56 | 0.75 | 0.55 | 0.56 | 2.34 | 0.62 | 0.29 | 0.60 | 0.78 | 0.70 | 0.56 | 2.25 |
| **RFR** | 0.67 | 0.44 | 0.48 | 0.69 | 0.83 | 0.54 | 1.86 | 0.73 | 0.53 | 0.40 | 0.63 | 0.76 | 0.50 | 1.42 |
| **Ridge** | 0.51 | 0.16 | 0.72 | 0.85 | 0.89 | 0.65 | 2.55 | 0.66 | 0.39 | 0.52 | 0.72 | 0.80 | 0.57 | 1.94 |
| **Lasso** | 0.00 | -0.01 | 0.87 | 0.93 | 1.40 | 0.75 | 1.97 | NaN | 0.00 | 0.85 | 0.92 | 1.18 | 0.74 | 2.21 |
| **GBR** | 0.64 | 0.40 | 0.51 | 0.72 | 0.89 | 0.58 | 1.74 | 0.71 | 0.50 | 0.43 | 0.65 | 0.77 | 0.52 | 1.60 |
| **MLPR** | 0.58 | 0.30 | 0.60 | 0.77 | 0.83 | 0.61 | 2.20 | 0.64 | 0.37 | 0.54 | 0.74 | 0.86 | 0.56 | 2.25 |
| **AdaBR** | 0.48 | 0.20 | 0.69 | 0.83 | 1.20 | 0.67 | 2.23 | 0.51 | 0.26 | 0.64 | 0.80 | 0.97 | 0.66 | 1.93 |
| **ENR** | 0.00 | -0.01 | 0.87 | 0.93 | 1.40 | 0.75 | 1.97 | NaN | 0.00 | 0.85 | 0.92 | 1.18 | 0.74 | 2.21 |
| **KRR** | 0.47 | 0.09 | 0.79 | 0.89 | 0.88 | 0.69 | 2.74 | 0.67 | 0.40 | 0.51 | 0.71 | 0.78 | 0.56 | 2.24 |
| **BRR** | 0.57 | 0.30 | 0.60 | 0.77 | 0.93 | 0.60 | 2.20 | 0.69 | 0.46 | 0.46 | 0.68 | 0.80 | 0.55 | 1.57 |

#R = Correlation coefficient, R² = Coefficient of determination, MSE = Mean squared error, RMSE = Root mean squared error, MAPE = Mean absolute percentage error, MAE = Mean absolute error, MRE = Maximum residual error, LR = Linear regression, SVR = Support vector regression, Ridge = Ridge regression, Lasso = =Lasso regression, GBR = Gradient Boosting regression, MLPR = MLP regression , AdaBR = AdaBoost regression, ENR = Elastic Net regression, KRR = Kernel Ridge regression and BRR = Bayesian Ridge regression

**Supplementary Table S5: Performance of Binary profile-based regressors for length group 11-20 residues on training and validation set**

| **Model** | **R** | **R2** | **MSE** | **RMSE** | **MAPE** | **MAE** | **MRE** | **R** | **R2** | **MSE** | **RMSE** | **MAPE** | **MAE** | **MRE** |
| --- | --- | --- | --- | --- | --- | --- | --- | --- | --- | --- | --- | --- | --- | --- |
|  | **TRAINING SET** | | | | | | | **VALIDATION SET** | | | | | | |
| **LR** | 0.35 | -0.89 | 1.38 | 1.17 | 3.90 | 0.88 | 3.76 | 0.37 | -0.99 | 1.42 | 1.19 | 18.46 | 0.87 | 5.28 |
| **SVR** | 0.72 | 0.51 | 0.35 | 0.59 | 2.81 | 0.46 | 2.20 | 0.75 | 0.56 | 0.31 | 0.56 | 19.97 | 0.43 | 1.74 |
| **DTR** | 0.59 | 0.21 | 0.57 | 0.76 | 1.45 | 0.55 | 2.55 | 0.67 | 0.32 | 0.48 | 0.69 | 7.05 | 0.51 | 2.29 |
| **RFR** | 0.70 | 0.49 | 0.37 | 0.61 | 2.23 | 0.48 | 1.98 | 0.74 | 0.55 | 0.32 | 0.57 | 21.77 | 0.44 | 1.92 |
| **Ridge** | 0.57 | 0.27 | 0.54 | 0.73 | 4.09 | 0.58 | 2.54 | 0.57 | 0.21 | 0.56 | 0.75 | 6.84 | 0.59 | 2.91 |
| **Lasso** | 0.00 | -0.01 | 0.73 | 0.86 | 5.05 | 0.68 | 2.80 | 0.00 | 0.00 | 0.71 | 0.84 | 40.65 | 0.67 | 3.40 |
| **GBR** | 0.68 | 0.45 | 0.40 | 0.64 | 3.51 | 0.52 | 1.91 | 0.71 | 0.49 | 0.37 | 0.61 | 23.95 | 0.49 | 1.79 |
| **MLPR** | 0.67 | 0.41 | 0.43 | 0.66 | 1.29 | 0.49 | 2.61 | 0.69 | 0.41 | 0.42 | 0.65 | 5.81 | 0.49 | 2.35 |
| **AdaBR** | 0.52 | 0.25 | 0.54 | 0.74 | 5.33 | 0.60 | 2.15 | 0.62 | 0.34 | 0.47 | 0.68 | 28.80 | 0.55 | 2.36 |
| **ENR** | 0.00 | -0.01 | 0.73 | 0.86 | 5.05 | 0.68 | 2.80 | 0.00 | 0.00 | 0.71 | 0.84 | 40.65 | 0.67 | 3.40 |
| **KRR** | 0.57 | 0.26 | 0.54 | 0.73 | 4.09 | 0.58 | 2.54 | 0.57 | 0.22 | 0.56 | 0.75 | 6.74 | 0.58 | 2.89 |
| **BRR** | 0.66 | 0.43 | 0.42 | 0.65 | 4.09 | 0.51 | 2.13 | 0.68 | 0.45 | 0.39 | 0.62 | 16.42 | 0.50 | 1.94 |

#R = Correlation coefficient, R² = Coefficient of determination, MSE = Mean squared error, RMSE = Root mean squared error, MAPE = Mean absolute percentage error, MAE = Mean absolute error, MRE = Maximum residual error, LR = Linear regression, SVR = Support vector regression, Ridge = Ridge regression, Lasso = =Lasso regression, GBR = Gradient Boosting regression, MLPR = MLP regression , AdaBR = AdaBoost regression, ENR = Elastic Net regression, KRR = Kernel Ridge regression and BRR = Bayesian Ridge regression

**Supplementary Table S6: Performance of Binary profile-based regressors for length group 21-40 residues on training and validation set**

| **Model** | **R** | **R2** | **MSE** | **RMSE** | **MAPE** | **MAE** | **MRE** | **R** | **R2** | **MSE** | **RMSE** | **MAPE** | **MAE** | **MRE** |
| --- | --- | --- | --- | --- | --- | --- | --- | --- | --- | --- | --- | --- | --- | --- |
|  | **TRAINING SET** | | | | | | | **VALIDATION SET** | | | | | | |
| **LR** | 0.01 | -8.86e+23 | 4.64e+23 | 6.81e+11 | 1.24e+11 | 8.29e+10 | 5.60e+12 | -0.05 | -6.84e+23 | 3.69e+23 | 6.08e+11 | 1.21e+11 | 6.60e+10 | 5.60e+12 |
| **SVR** | 0.65 | 0.41 | 0.31 | 0.55 | 48.64 | 0.44 | 1.74 | 0.56 | 0.31 | 0.37 | 0.61 | 29.67 | 0.47 | 2.99 |
| **DTR** | 0.46 | -0.02 | 0.54 | 0.73 | 23.87 | 0.55 | 2.48 | 0.28 | -0.43 | 0.77 | 0.88 | 35.40 | 0.64 | 3.03 |
| **RFR** | 0.60 | 0.36 | 0.33 | 0.58 | 39.81 | 0.46 | 1.69 | 0.53 | 0.28 | 0.39 | 0.62 | 28.25 | 0.49 | 2.46 |
| **Ridge** | 0.50 | -0.12 | 0.58 | 0.76 | 43.43 | 0.58 | 2.54 | 0.34 | -0.49 | 0.80 | 0.90 | 33.02 | 0.66 | 4.46 |
| **Lasso** | 0.00 | 0.00 | 0.53 | 0.73 | 39.54 | 0.60 | 1.89 | 0.00 | 0.00 | 0.54 | 0.73 | 33.80 | 0.60 | 2.59 |
| **GBR** | 0.59 | 0.33 | 0.35 | 0.59 | 40.42 | 0.48 | 2.05 | 0.46 | 0.21 | 0.43 | 0.65 | 26.88 | 0.52 | 2.94 |
| **MLPR** | 0.59 | 0.29 | 0.37 | 0.61 | 36.75 | 0.46 | 2.09 | 0.45 | 0.03 | 0.53 | 0.72 | 27.91 | 0.53 | 3.85 |
| **AdaBR** | 0.37 | 0.08 | 0.48 | 0.69 | 41.77 | 0.58 | 1.94 | 0.23 | 0.05 | 0.52 | 0.72 | 33.80 | 0.59 | 2.73 |
| **ENR** | 0.00 | 0.00 | 0.53 | 0.73 | 39.54 | 0.60 | 1.89 | 0.00 | 0.00 | 0.54 | 0.73 | 33.80 | 0.60 | 2.59 |
| **KRR** | 0.50 | -0.12 | 0.58 | 0.76 | 43.32 | 0.58 | 2.52 | 0.34 | -0.49 | 0.81 | 0.90 | 33.22 | 0.66 | 4.47 |
| **BRR** | 0.59 | 0.35 | 0.34 | 0.58 | 50.34 | 0.46 | 1.97 | 0.44 | 0.17 | 0.45 | 0.67 | 27.83 | 0.52 | 3.35 |

#R = Correlation coefficient, R² = Coefficient of determination, MSE = Mean squared error, RMSE = Root mean squared error, MAPE = Mean absolute percentage error, MAE = Mean absolute error, MRE = Maximum residual error, LR = Linear regression, SVR = Support vector regression, Ridge = Ridge regression, Lasso = =Lasso regression, GBR = Gradient Boosting regression, MLPR = MLP regression , AdaBR = AdaBoost regression, ENR = Elastic Net regression, KRR = Kernel Ridge regression and BRR = Bayesian Ridge regression

**Supplementary Table S7: Performance of Binary profile-based regressors for length group 41 or more residues on training and validation set**

| **Model** | **R** | **R2** | **MSE** | **RMSE** | **MAPE** | **MAE** | **MRE** | **R** | **R2** | **MSE** | **RMSE** | **MAPE** | **MAE** | **MRE** |
| --- | --- | --- | --- | --- | --- | --- | --- | --- | --- | --- | --- | --- | --- | --- |
|  | **TRAINING SET** | | | | | | | **VALIDATION SET** | | | | | | |
| **LR** | 0.51 | 0.18 | 0.47 | 0.68 | 1.75 | 0.52 | 1.64 | 0.42 | 0.13 | 0.52 | 0.72 | 3.83 | 0.54 | 2.16 |
| **SVR** | 0.52 | 0.18 | 0.47 | 0.69 | 1.73 | 0.52 | 1.63 | 0.44 | 0.18 | 0.49 | 0.70 | 6.64 | 0.55 | 1.70 |
| **DTR** | 0.15 | -1.01 | 1.15 | 1.07 | 2.58 | 0.78 | 2.72 | 0.26 | -0.33 | 0.79 | 0.89 | 2.09 | 0.70 | 2.06 |
| **RFR** | 0.55 | 0.21 | 0.45 | 0.67 | 1.65 | 0.52 | 1.50 | 0.48 | 0.21 | 0.47 | 0.69 | 10.22 | 0.53 | 1.87 |
| **Ridge** | 0.53 | 0.21 | 0.45 | 0.67 | 1.73 | 0.50 | 1.58 | 0.43 | 0.16 | 0.50 | 0.71 | 3.19 | 0.54 | 2.08 |
| **Lasso** | NaN | -0.05 | 0.60 | 0.77 | 1.80 | 0.63 | 1.59 | 0.00 | 0.00 | 0.60 | 0.77 | 36.30 | 0.64 | 1.80 |
| **GBR** | 0.33 | 0.00 | 0.57 | 0.76 | 1.74 | 0.60 | 1.91 | 0.26 | -0.02 | 0.61 | 0.78 | 6.96 | 0.57 | 2.57 |
| **MLPR** | 0.53 | 0.27 | 0.41 | 0.64 | 1.53 | 0.49 | 1.41 | 0.42 | 0.15 | 0.51 | 0.71 | 5.47 | 0.54 | 2.25 |
| **AdaBR** | 0.28 | -0.01 | 0.58 | 0.76 | 1.72 | 0.62 | 1.96 | 0.39 | 0.14 | 0.51 | 0.72 | 18.91 | 0.57 | 1.53 |
| **ENR** | NaN | -0.05 | 0.60 | 0.77 | 1.80 | 0.63 | 1.59 | 0.00 | 0.00 | 0.60 | 0.77 | 36.30 | 0.64 | 1.80 |
| **KRR** | 0.49 | 0.21 | 0.45 | 0.67 | 1.67 | 0.51 | 1.69 | 0.39 | 0.10 | 0.54 | 0.73 | 3.65 | 0.56 | 2.28 |
| **BRR** | 0.55 | 0.24 | 0.43 | 0.66 | 1.71 | 0.50 | 1.52 | 0.46 | 0.20 | 0.48 | 0.69 | 4.41 | 0.53 | 1.90 |

#R = Correlation coefficient, R² = Coefficient of determination, MSE = Mean squared error, RMSE = Root mean squared error, MAPE = Mean absolute percentage error, MAE = Mean absolute error, MRE = Maximum residual error, LR = Linear regression, SVR = Support vector regression, Ridge = Ridge regression, Lasso = =Lasso regression, GBR = Gradient Boosting regression, MLPR = MLP regression , AdaBR = AdaBoost regression, ENR = Elastic Net regression, KRR = Kernel Ridge regression and BRR = Bayesian Ridge regression

**Supplementary Table S8: Performance of ProtBERT embeddings-based regressors on training and validation set**

| **Model** | **R** | **R2** | **MSE** | **RMSE** | **MAPE** | **MAE** | **MRE** | **R** | **R2** | **MSE** | **RMSE** | **MAPE** | **MAE** | **MRE** |
| --- | --- | --- | --- | --- | --- | --- | --- | --- | --- | --- | --- | --- | --- | --- |
|  | **TRAINING SET** | | | | | | | **VALIDATION SET** | | | | | | |
| **LR** | 0.47 | -0.13 | 0.77 | 0.88 | 65.81 | 0.65 | 5.10 | 0.57 | 0.11 | 0.61 | 0.78 | 93.92 | 0.60 | 3.52 |
| **SVR** | 0.54 | 0.28 | 0.49 | 0.70 | 98.41 | 0.55 | 2.61 | 0.60 | 0.34 | 0.45 | 0.67 | 80.14 | 0.53 | 2.61 |
| **DTR** | 0.36 | -0.31 | 0.89 | 0.94 | 62.02 | 0.71 | 3.70 | 0.39 | -0.18 | 0.81 | 0.90 | 33.81 | 0.68 | 3.85 |
| **RFR** | **0.65** | **0.40** | **0.41** | **0.64** | **103.72** | **0.51** | **2.39** | **0.70** | **0.46** | **0.37** | **0.61** | **110.53** | **0.49** | **2.45** |
| **Ridge** | 0.60 | 0.36 | 0.44 | 0.66 | 79.10 | 0.53 | 2.42 | 0.67 | 0.44 | 0.38 | 0.62 | 86.96 | 0.49 | 2.41 |
| **Lasso** | 0.00 | -0.01 | 0.69 | 0.83 | 131.08 | 0.67 | 2.87 | NAN | 0.00 | 0.69 | 0.83 | 172.03 | 0.67 | 2.96 |
| **GBR** | 0.61 | 0.37 | 0.43 | 0.66 | 97.87 | 0.52 | 2.35 | 0.67 | 0.45 | 0.38 | 0.62 | 97.77 | 0.49 | 2.40 |
| **MLPR** | 0.59 | 0.24 | 0.52 | 0.72 | 115.12 | 0.55 | 2.79 | 0.67 | 0.38 | 0.43 | 0.66 | 54.89 | 0.51 | 2.79 |
| **AdaBR** | 0.54 | 0.27 | 0.50 | 0.71 | 94.49 | 0.58 | 2.44 | 0.58 | 0.30 | 0.48 | 0.69 | 131.60 | 0.56 | 2.72 |
| **ENR** | 0.00 | -0.01 | 0.69 | 0.83 | 131.08 | 0.67 | 2.87 | NAN | 0.00 | 0.69 | 0.83 | 172.03 | 0.67 | 2.96 |
| **KRR** | 0.60 | 0.36 | 0.44 | 0.66 | 79.77 | 0.53 | 2.42 | 0.67 | 0.44 | 0.38 | 0.62 | 86.78 | 0.49 | 2.40 |
| **BRR** | 0.61 | 0.37 | 0.43 | 0.66 | 75.92 | 0.52 | 2.40 | 0.68 | 0.45 | 0.38 | 0.61 | 92.35 | 0.49 | 2.33 |

#R = Correlation coefficient, R² = Coefficient of determination, MSE = Mean squared error, RMSE = Root mean squared error, MAPE = Mean absolute percentage error, MAE = Mean absolute error, MRE = Maximum residual error, LR = Linear regression, SVR = Support vector regression, Ridge = Ridge regression, Lasso = =Lasso regression, GBR = Gradient Boosting regression, MLPR = MLP regression , AdaBR = AdaBoost regression, ENR = Elastic Net regression, KRR = Kernel Ridge regression and BRR = Bayesian Ridge regression

**Supplementary Table S9: Performance of BioBERT embeddings-based regressors on training and validation set**

| **Model** | **R** | **R2** | **MSE** | **RMSE** | **MAPE** | **MAE** | **MRE** | **R** | **R2** | **MSE** | **RMSE** | **MAPE** | **MAE** | **MRE** |
| --- | --- | --- | --- | --- | --- | --- | --- | --- | --- | --- | --- | --- | --- | --- |
|  | **TRAINING SET** | | | | | | | **VALIDATION SET** | | | | | | |
| **LR** | 0.50 | 0.12 | 0.60 | 0.78 | 57.42 | 0.61 | 3.00 | 0.55 | 0.19 | 0.56 | 0.75 | 93.66 | 0.59 | 2.40 |
| **SVR** | 0.52 | 0.26 | 0.51 | 0.71 | 107.31 | 0.57 | 2.51 | 0.59 | 0.32 | 0.47 | 0.68 | 111.91 | 0.54 | 2.52 |
| **DTR** | 0.28 | -0.49 | 1.02 | 1.01 | 195.49 | 0.78 | 3.58 | 0.33 | -0.42 | 0.98 | 0.99 | 92.73 | 0.75 | 3.75 |
| **RFR** | **0.58** | **0.31** | **0.47** | **0.69** | **93.58** | **0.55** | **2.45** | **0.63** | **0.36** | **0.44** | **0.66** | **75.33** | **0.53** | **2.64** |
| **Ridge** | 0.55 | 0.28 | 0.49 | 0.70 | 71.42 | 0.56 | 2.43 | 0.62 | 0.36 | 0.44 | 0.66 | 94.40 | 0.53 | 2.40 |
| **Lasso** | 0.00 | -0.01 | 0.69 | 0.83 | 131.08 | 0.67 | 2.87 | NAN | 0.00 | 0.69 | 0.83 | 172.03 | 0.67 | 2.96 |
| **GBR** | 0.56 | 0.30 | 0.48 | 0.69 | 98.29 | 0.55 | 2.45 | 0.60 | 0.35 | 0.45 | 0.67 | 90.87 | 0.54 | 2.43 |
| **MLPR** | 0.53 | 0.15 | 0.58 | 0.76 | 87.62 | 0.59 | 2.94 | 0.63 | 0.30 | 0.48 | 0.69 | 55.28 | 0.54 | 2.27 |
| **AdaBR** | 0.47 | 0.21 | 0.54 | 0.74 | 99.48 | 0.60 | 2.38 | 0.52 | 0.26 | 0.51 | 0.71 | 128.14 | 0.59 | 2.51 |
| **ENR** | 0.00 | -0.01 | 0.69 | 0.83 | 131.08 | 0.67 | 2.87 | NAN | 0.00 | 0.69 | 0.83 | 172.03 | 0.67 | 2.96 |
| **KRR** | 0.55 | 0.28 | 0.49 | 0.70 | 72.03 | 0.56 | 2.44 | 0.62 | 0.37 | 0.44 | 0.66 | 93.76 | 0.53 | 2.39 |
| **BRR** | 0.56 | 0.31 | 0.47 | 0.69 | 84.11 | 0.55 | 2.26 | 0.63 | 0.39 | 0.42 | 0.65 | 97.54 | 0.52 | 2.41 |

#R = Correlation coefficient, R² = Coefficient of determination, MSE = Mean squared error, RMSE = Root mean squared error, MAPE = Mean absolute percentage error, MAE = Mean absolute error, MRE = Maximum residual error, LR = Linear regression, SVR = Support vector regression, Ridge = Ridge regression, Lasso = =Lasso regression, GBR = Gradient Boosting regression, MLPR = MLP regression , AdaBR = AdaBoost regression, ENR = Elastic Net regression, KRR = Kernel Ridge regression and BRR = Bayesian Ridge regression

**Supplementary Table S10: Performance of ESM-2 embeddings-based regressors on training and validation set**

| **Model** | **R** | **R2** | **MSE** | **RMSE** | **MAPE** | **MAE** | **MRE** | **R** | **R2** | **MSE** | **RMSE** | **MAPE** | **MAE** | **MRE** |
| --- | --- | --- | --- | --- | --- | --- | --- | --- | --- | --- | --- | --- | --- | --- |
|  | **TRAINING SET** | | | | | | | **VALIDATION SET** | | | | | | |
| **LR** | 0.45 | -0.40 | 0.95 | 0.97 | 78.18 | 0.69 | 7.64 | 0.52 | -0.04 | 0.72 | 0.85 | 242.83 | 0.64 | 5.66 |
| **SVR** | 0.61 | 0.37 | 0.43 | 0.66 | 97.97 | 0.51 | 2.68 | 0.66 | 0.43 | 0.39 | 0.63 | 159.50 | 0.50 | 2.53 |
| **DTR** | 0.42 | -0.18 | 0.80 | 0.89 | 113.62 | 0.67 | 3.22 | 0.49 | -0.03 | 0.71 | 0.84 | 283.39 | 0.63 | 3.04 |
| **RFR** | **0.69** | **0.46** | **0.37** | **0.61** | **81.88** | **0.48** | **2.35** | **0.76** | **0.55** | **0.31** | **0.56** | **150.85** | **0.44** | **2.43** |
| **Ridge** | 0.66 | 0.43 | 0.39 | 0.63 | 76.46 | 0.49 | 2.71 | 0.71 | 0.50 | 0.34 | 0.59 | 219.12 | 0.46 | 2.24 |
| **Lasso** | 0.00 | -0.01 | 0.69 | 0.83 | 131.08 | 0.67 | 2.87 | NAN | 0.00 | 0.69 | 0.83 | 172.03 | 0.67 | 2.96 |
| **GBR** | 0.65 | 0.42 | 0.39 | 0.63 | 65.80 | 0.50 | 2.37 | 0.72 | 0.51 | 0.33 | 0.58 | 191.35 | 0.46 | 2.16 |
| **MLPR** | 0.63 | 0.31 | 0.47 | 0.69 | 86.09 | 0.51 | 3.07 | 0.72 | 0.48 | 0.36 | 0.60 | 300.35 | 0.46 | 2.41 |
| **AdaBR** | 0.59 | 0.33 | 0.45 | 0.67 | 98.25 | 0.55 | 2.40 | 0.64 | 0.38 | 0.42 | 0.65 | 182.88 | 0.53 | 2.33 |
| **ENR** | 0.00 | -0.01 | 0.69 | 0.83 | 131.08 | 0.67 | 2.87 | NAN | 0.00 | 0.69 | 0.83 | 172.03 | 0.67 | 2.96 |
| **KRR** | 0.66 | 0.43 | 0.39 | 0.63 | 75.80 | 0.49 | 2.68 | 0.71 | 0.50 | 0.34 | 0.59 | 217.43 | 0.46 | 2.24 |
| **BRR** | 0.66 | 0.43 | 0.39 | 0.63 | 80.80 | 0.49 | 2.61 | 0.71 | 0.50 | 0.35 | 0.59 | 199.21 | 0.47 | 2.30 |

#R = Correlation coefficient, R² = Coefficient of determination, MSE = Mean squared error, RMSE = Root mean squared error, MAPE = Mean absolute percentage error, MAE = Mean absolute error, MRE = Maximum residual error, LR = Linear regression, SVR = Support vector regression, Ridge = Ridge regression, Lasso = =Lasso regression, GBR = Gradient Boosting regression, MLPR = MLP regression , AdaBR = AdaBoost regression, ENR = Elastic Net regression, KRR = Kernel Ridge regression and BRR = Bayesian Ridge regression

**Supplementary Table S11: Performance of RF-based regressor on Top 200 selected features on training and validation set**

| **Model** | **R** | **R2** | **MSE** | **RMSE** | **MAPE** | **MAE** | **MRE** | **R** | **R2** | **MSE** | **RMSE** | **MAPE** | **MAE** | **MRE** |
| --- | --- | --- | --- | --- | --- | --- | --- | --- | --- | --- | --- | --- | --- | --- |
|  | **TRAINING SET** | | | | | | | **VALIDATION SET** | | | | | | |
| **LR** | 0.62 | 0.38 | 0.42 | 0.65 | 97.20 | 0.51 | 2.49 | 0.08 | -1307.78 | 897.91 | 29.97 | 130.14 | 1.58 | 839.84 |
| **SVR** | 0.51 | 0.26 | 0.50 | 0.71 | 189.37 | 0.57 | 2.45 | 0.59 | 0.33 | 0.46 | 0.68 | 131.30 | 0.54 | 2.60 |
| **DTR** | 0.46 | -0.07 | 0.73 | 0.85 | 103.43 | 0.64 | 3.00 | 0.58 | 0.15 | 0.58 | 0.76 | 264.25 | 0.55 | 2.85 |
| **RFR** | **0.71** | **0.50** | **0.34** | **0.58** | **143.45** | **0.46** | **2.61** | **0.76** | **0.57** | **0.30** | **0.55** | **107.74** | **0.42** | **2.06** |
| **Ridge** | 0.63 | 0.39 | 0.42 | 0.64 | 105.45 | 0.51 | 2.51 | 0.59 | 0.33 | 0.46 | 0.68 | 123.35 | 0.51 | 5.75 |
| **Lasso** | 0.37 | 0.14 | 0.59 | 0.77 | 204.30 | 0.62 | 2.46 | 0.47 | 0.21 | 0.54 | 0.74 | 157.28 | 0.58 | 2.80 |
| **GBR** | 0.65 | 0.42 | 0.40 | 0.63 | 155.59 | 0.50 | 2.54 | 0.71 | 0.49 | 0.35 | 0.59 | 110.78 | 0.47 | 2.28 |
| **MLPR** | 0.60 | 0.31 | 0.47 | 0.69 | 244.00 | 0.54 | 2.61 | 0.55 | 0.22 | 0.53 | 0.73 | 190.25 | 0.57 | 3.87 |
| **AdaBR** | 0.56 | 0.30 | 0.48 | 0.69 | 169.83 | 0.56 | 2.26 | 0.62 | 0.36 | 0.44 | 0.66 | 115.98 | 0.54 | 2.42 |
| **ENR** | 0.41 | 0.17 | 0.57 | 0.75 | 208.50 | 0.60 | 2.48 | 0.50 | 0.24 | 0.52 | 0.72 | 154.95 | 0.57 | 2.78 |
| **KRR** | 0.62 | 0.38 | 0.42 | 0.65 | 56.56 | 0.51 | 2.50 | 0.57 | 0.30 | 0.48 | 0.69 | 128.49 | 0.52 | 5.72 |
| **BRR** | 0.63 | 0.39 | 0.41 | 0.64 | 176.00 | 0.51 | 2.31 | 0.66 | 0.44 | 0.39 | 0.62 | 141.95 | 0.50 | 2.13 |

#R = Correlation coefficient, R² = Coefficient of determination, MSE = Mean squared error, RMSE = Root mean squared error, MAPE = Mean absolute percentage error, MAE = Mean absolute error, MRE = Maximum residual error, LR = Linear regression, SVR = Support vector regression, Ridge = Ridge regression, Lasso = =Lasso regression, GBR = Gradient Boosting regression, MLPR = MLP regression , AdaBR = AdaBoost regression, ENR = Elastic Net regression, KRR = Kernel Ridge regression and BRR = Bayesian Ridge regression

**Supplementary Table S12: Performance of RF-based regressor on Top 500 selected features on training and validation set**

| **Model** | **R** | **R2** | **MSE** | **RMSE** | **MAPE** | **MAE** | **MRE** | **R** | **R2** | **MSE** | **RMSE** | **MAPE** | **MAE** | **MRE** |
| --- | --- | --- | --- | --- | --- | --- | --- | --- | --- | --- | --- | --- | --- | --- |
|  | **TRAINING SET** | | | | | | | **VALIDATION SET** | | | | | | |
| **LR** | 0.66 | 0.36 | 0.44 | 0.66 | 75.52 | 0.48 | 5.68 | -0.04 | -302.19 | 208.01 | 14.42 | 220.77 | 1.01 | 403.86 |
| **SVR** | 0.52 | 0.27 | 0.50 | 0.70 | 194.38 | 0.56 | 2.43 | 0.60 | 0.35 | 0.45 | 0.67 | 127.08 | 0.53 | 2.61 |
| **DTR** | 0.54 | 0.07 | 0.63 | 0.80 | 144.29 | 0.59 | 3.21 | 0.61 | 0.17 | 0.57 | 0.76 | 28.97 | 0.54 | 3.08 |
| **RFR** | 0.72 | 0.52 | 0.33 | 0.57 | 124.69 | 0.44 | 2.47 | 0.77 | 0.58 | 0.29 | 0.53 | 95.51 | 0.41 | 2.03 |
| **Ridge** | 0.66 | 0.38 | 0.42 | 0.65 | 68.72 | 0.48 | 5.55 | 0.63 | 0.37 | 0.43 | 0.66 | 221.47 | 0.49 | 3.51 |
| **Lasso** | 0.38 | 0.14 | 0.59 | 0.76 | 203.73 | 0.61 | 2.46 | 0.48 | 0.21 | 0.54 | 0.73 | 156.18 | 0.58 | 2.81 |
| **GBR** | 0.66 | 0.43 | 0.38 | 0.62 | 131.02 | 0.49 | 2.49 | 0.72 | 0.50 | 0.34 | 0.58 | 110.60 | 0.46 | 2.27 |
| **MLPR** | 0.64 | 0.39 | 0.42 | 0.64 | 39.77 | 0.50 | 3.00 | 0.59 | 0.31 | 0.48 | 0.69 | 162.11 | 0.52 | 4.05 |
| **AdaBR** | 0.58 | 0.33 | 0.46 | 0.68 | 167.84 | 0.55 | 2.13 | 0.63 | 0.38 | 0.43 | 0.65 | 111.02 | 0.54 | 2.09 |
| **ENR** | 0.42 | 0.17 | 0.56 | 0.75 | 200.75 | 0.60 | 2.44 | 0.51 | 0.25 | 0.52 | 0.72 | 147.29 | 0.57 | 2.81 |
| **KRR** | 0.66 | 0.37 | 0.43 | 0.65 | 72.75 | 0.48 | 5.53 | 0.63 | 0.37 | 0.43 | 0.66 | 219.96 | 0.49 | 3.54 |
| **BRR** | 0.68 | 0.47 | 0.36 | 0.60 | 52.99 | 0.48 | 2.33 | 0.70 | 0.49 | 0.35 | 0.59 | 168.80 | 0.47 | 1.97 |

#R = Correlation coefficient, R² = Coefficient of determination, MSE = Mean squared error, RMSE = Root mean squared error, MAPE = Mean absolute percentage error, MAE = Mean absolute error, MRE = Maximum residual error, LR = Linear regression, SVR = Support vector regression, Ridge = Ridge regression, Lasso = =Lasso regression, GBR = Gradient Boosting regression, MLPR = MLP regression , AdaBR = AdaBoost regression, ENR = Elastic Net regression, KRR = Kernel Ridge regression and BRR = Bayesian Ridge regression

**Supplementary Table S13: Performance of RF-based regressor on Top 1000 selected features on training and validation set**

| **Model** | **R** | **R2** | **MSE** | **RMSE** | **MAPE** | **MAE** | **MRE** | **R** | **R2** | **MSE** | **RMSE** | **MAPE** | **MAE** | **MRE** |
| --- | --- | --- | --- | --- | --- | --- | --- | --- | --- | --- | --- | --- | --- | --- |
|  | **TRAINING SET** | | | | | | | **VALIDATION SET** | | | | | | |
| **LR** | 0.62 | 0.22 | 0.53 | 0.73 | 105.93 | 0.52 | 4.60 | -0.05 | -827.72 | 568.56 | 23.84 | 263.73 | 1.42 | 668.10 |
| **SVR** | 0.39 | 0.15 | 0.58 | 0.76 | 174.12 | 0.61 | 2.55 | 0.40 | 0.16 | 0.58 | 0.76 | 72.94 | 0.61 | 3.06 |
| **DTR** | 0.52 | 0.06 | 0.64 | 0.80 | 15.65 | 0.59 | 2.90 | 0.57 | 0.14 | 0.59 | 0.77 | 209.51 | 0.56 | 2.81 |
| **RFR** | **0.73** | **0.52** | **0.33** | **0.57** | **111.29** | **0.44** | **2.58** | **0.78** | **0.59** | **0.28** | **0.53** | **89.25** | **0.41** | **2.12** |
| **Ridge** | 0.66 | 0.34 | 0.45 | 0.67 | 91.33 | 0.49 | 4.28 | 0.54 | 0.06 | 0.64 | 0.80 | 58.63 | 0.54 | 10.02 |
| **Lasso** | 0.38 | 0.14 | 0.58 | 0.76 | 222.28 | 0.61 | 2.54 | 0.49 | 0.23 | 0.53 | 0.73 | 127.80 | 0.58 | 2.84 |
| **GBR** | 0.67 | 0.44 | 0.38 | 0.62 | 128.65 | 0.49 | 2.48 | 0.72 | 0.50 | 0.34 | 0.59 | 93.05 | 0.47 | 2.25 |
| **MLPR** | 0.62 | 0.01 | 0.68 | 0.82 | 46.63 | 0.64 | 2.94 | 0.52 | -0.24 | 0.85 | 0.92 | 347.93 | 0.72 | 2.70 |
| **AdaBR** | 0.59 | 0.33 | 0.46 | 0.68 | 174.96 | 0.55 | 2.27 | 0.64 | 0.37 | 0.43 | 0.66 | 101.79 | 0.54 | 2.09 |
| **ENR** | 0.42 | 0.18 | 0.56 | 0.75 | 234.28 | 0.60 | 2.56 | 0.52 | 0.26 | 0.51 | 0.71 | 114.95 | 0.57 | 2.82 |
| **KRR** | 0.66 | 0.34 | 0.45 | 0.67 | 91.33 | 0.49 | 4.28 | 0.54 | 0.06 | 0.64 | 0.80 | 58.60 | 0.54 | 10.02 |
| **BRR** | 0.70 | 0.49 | 0.35 | 0.59 | 40.76 | 0.47 | 2.45 | 0.70 | 0.49 | 0.35 | 0.59 | 149.07 | 0.47 | 2.12 |

#R = Correlation coefficient, R² = Coefficient of determination, MSE = Mean squared error, RMSE = Root mean squared error, MAPE = Mean absolute percentage error, MAE = Mean absolute error, MRE = Maximum residual error, LR = Linear regression, SVR = Support vector regression, Ridge = Ridge regression, Lasso = =Lasso regression, GBR = Gradient Boosting regression, MLPR = MLP regression , AdaBR = AdaBoost regression, ENR = Elastic Net regression, KRR = Kernel Ridge regression and BRR = Bayesian Ridge regression

**Supplementary Table S14: Performance of RF-based regressor on Top 1500 selected features on training and validation set**

| **Model** | **R** | **R2** | **MSE** | **RMSE** | **MAPE** | **MAE** | **MRE** | **R** | **R2** | **MSE** | **RMSE** | **MAPE** | **MAE** | **MRE** |
| --- | --- | --- | --- | --- | --- | --- | --- | --- | --- | --- | --- | --- | --- | --- |
|  | **TRAINING SET** | | | | | | | **VALIDATION SET** | | | | | | |
| **LR** | 0.30 | -1.84 | 1.93 | 1.39 | 117.43 | 0.72 | 22.92 | 0.17 | -4.56 | 3.82 | 1.95 | 51.12 | 0.75 | 43.47 |
| **SVR** | 0.39 | 0.15 | 0.58 | 0.76 | 175.23 | 0.61 | 2.55 | 0.40 | 0.16 | 0.58 | 0.76 | 73.10 | 0.61 | 3.06 |
| **DTR** | 0.52 | 0.00 | 0.68 | 0.83 | 18.67 | 0.60 | 3.51 | 0.63 | 0.24 | 0.52 | 0.72 | 117.05 | 0.52 | 2.92 |
| **RFR** | **0.72** | **0.52** | **0.33** | **0.57** | **115.00** | **0.44** | **2.47** | **0.77** | **0.59** | **0.28** | **0.53** | **92.93** | **0.41** | **2.12** |
| **Ridge** | 0.56 | 0.03 | 0.66 | 0.81 | 113.27 | 0.56 | 7.09 | 0.53 | 0.05 | 0.65 | 0.81 | 248.81 | 0.58 | 5.42 |
| **Lasso** | 0.38 | 0.14 | 0.58 | 0.76 | 222.28 | 0.61 | 2.54 | 0.49 | 0.23 | 0.53 | 0.73 | 127.80 | 0.58 | 2.84 |
| **GBR** | 0.66 | 0.43 | 0.39 | 0.62 | 94.10 | 0.50 | 2.35 | 0.72 | 0.51 | 0.34 | 0.58 | 103.52 | 0.46 | 2.29 |
| **MLPR** | 0.64 | 0.19 | 0.55 | 0.74 | 56.77 | 0.59 | 3.62 | 0.57 | 0.09 | 0.63 | 0.79 | 96.73 | 0.61 | 3.48 |
| **AdaBR** | 0.61 | 0.35 | 0.44 | 0.67 | 159.90 | 0.54 | 2.34 | 0.64 | 0.38 | 0.42 | 0.65 | 122.23 | 0.53 | 1.98 |
| **ENR** | 0.42 | 0.18 | 0.56 | 0.75 | 234.28 | 0.60 | 2.56 | 0.52 | 0.26 | 0.51 | 0.71 | 114.95 | 0.57 | 2.82 |
| **KRR** | 0.56 | 0.03 | 0.66 | 0.81 | 113.23 | 0.56 | 7.09 | 0.53 | 0.05 | 0.65 | 0.81 | 248.83 | 0.58 | 5.42 |
| **BRR** | 0.69 | 0.48 | 0.35 | 0.59 | 31.08 | 0.47 | 2.53 | 0.70 | 0.49 | 0.35 | 0.59 | 165.53 | 0.47 | 2.13 |

#R = Correlation coefficient, R² = Coefficient of determination, MSE = Mean squared error, RMSE = Root mean squared error, MAPE = Mean absolute percentage error, MAE = Mean absolute error, MRE = Maximum residual error, LR = Linear regression, SVR = Support vector regression, Ridge = Ridge regression, Lasso = =Lasso regression, GBR = Gradient Boosting regression, MLPR = MLP regression , AdaBR = AdaBoost regression, ENR = Elastic Net regression, KRR = Kernel Ridge regression and BRR = Bayesian Ridge regression

**Supplementary Table S15: Performance of RF-based regressor on Top 2000 selected features on training and validation set**

| **Model** | **R** | **R2** | **MSE** | **RMSE** | **MAPE** | **MAE** | **MRE** | **R** | **R2** | **MSE** | **RMSE** | **MAPE** | **MAE** | **MRE** |
| --- | --- | --- | --- | --- | --- | --- | --- | --- | --- | --- | --- | --- | --- | --- |
|  | **TRAINING SET** | | | | | | | **VALIDATION SET** | | | | | | |
| **LR** | 0.16 | -8.00 | 6.12 | 2.47 | 84.34 | 1.16 | 29.88 | -0.07 | -33452.8 | 22951.6 | 151.50 | 1516.0 | 6.55 | 4246.8 |
| **SVR** | 0.39 | 0.15 | 0.58 | 0.76 | 175.71 | 0.61 | 2.55 | 0.40 | 0.16 | 0.58 | 0.76 | 73.13 | 0.61 | 3.06 |
| **DTR** | 0.52 | 0.02 | 0.66 | 0.82 | 23.90 | 0.60 | 3.68 | 0.60 | 0.20 | 0.55 | 0.74 | 116.37 | 0.53 | 3.06 |
| **RFR** | **0.72** | **0.52** | **0.33** | **0.57** | **117.67** | **0.44** | **2.39** | **0.77** | **0.59** | **0.28** | **0.53** | **90.83** | **0.41** | **2.06** |
| **Ridge** | 0.54 | -0.13 | 0.77 | 0.88 | 79.99 | 0.62 | 5.06 | 0.48 | -0.23 | 0.85 | 0.92 | 389.96 | 0.65 | 5.30 |
| **Lasso** | 0.38 | 0.14 | 0.58 | 0.76 | 222.28 | 0.61 | 2.54 | 0.49 | 0.23 | 0.53 | 0.73 | 127.80 | 0.58 | 2.84 |
| **GBR** | 0.66 | 0.43 | 0.38 | 0.62 | 109.27 | 0.49 | 2.35 | 0.73 | 0.51 | 0.34 | 0.58 | 116.28 | 0.46 | 2.26 |
| **MLPR** | 0.61 | 0.07 | 0.63 | 0.79 | 24.33 | 0.62 | 3.66 | 0.50 | -0.32 | 0.91 | 0.95 | 275.77 | 0.66 | 11.44 |
| **AdaBR** | 0.58 | 0.32 | 0.47 | 0.68 | 144.89 | 0.55 | 2.34 | 0.63 | 0.37 | 0.43 | 0.66 | 114.28 | 0.54 | 2.11 |
| **ENR** | 0.42 | 0.18 | 0.56 | 0.75 | 234.28 | 0.60 | 2.56 | 0.52 | 0.26 | 0.51 | 0.71 | 114.95 | 0.57 | 2.82 |
| **KRR** | 0.54 | -0.13 | 0.77 | 0.88 | 79.99 | 0.62 | 5.06 | 0.48 | -0.23 | 0.85 | 0.92 | 389.96 | 0.65 | 5.30 |
| **BRR** | 0.69 | 0.48 | 0.36 | 0.60 | 18.21 | 0.48 | 2.44 | 0.71 | 0.50 | 0.34 | 0.59 | 167.90 | 0.47 | 2.19 |

#R = Correlation coefficient, R² = Coefficient of determination, MSE = Mean squared error, RMSE = Root mean squared error, MAPE = Mean absolute percentage error, MAE = Mean absolute error, MRE = Maximum residual error, LR = Linear regression, SVR = Support vector regression, Ridge = Ridge regression, Lasso = =Lasso regression, GBR = Gradient Boosting regression, MLPR = MLP regression , AdaBR = AdaBoost regression, ENR = Elastic Net regression, KRR = Kernel Ridge regression and BRR = Bayesian Ridge regression
